# The life history traits of phages in a cocktail determine coinfection dynamics and efficacy

**DOI:** 10.1101/2024.10.26.620386

**Authors:** Ruqaiyah Khan, Harshali V. Chaudhari, Mandar M. Inamdar, Kiran Kondabagil

## Abstract

Phage cocktails are preferred over single phages for efficacious and broader-spectrum therapy. An ideal phage cocktail should have a minimum number of phages with efficient infection kinetics and delay the emergence of resistance in bacterial populations. This study examined population dynamics of the common host and combinations of two phages (N4, KKE5P, and Ec_YwIITB1) through experimental and modeling approaches to gaining insights into how phage life history traits influence the outcome of infection in the short-term of approximately an infection cycle, and whether it is informative for developing efficacious cocktails. We tested the killing efficacy of a cocktail containing two divergent phages (N4 and Ec_YwIITB1) with similar adsorption rates but differed in their latency period. Because of the shorter latency period, phage N4 dominated under all conditions tested. The cocktail essentially behaves as a single phage. When two phages (N4 and KKE5P), with similar adsorption rates and latency periods but targeted different host receptors were used, it not only resulted in the efficient replication of both phages but also improved the suppression of the emergence of resistance when compared to the N4 and Ec_YwIITB1 combination, thus behaving like an ideal cocktail. The ODE-based mathematical model demonstrated predictive capabilities consistent with experimental observations and offered insights into infection dynamics. The model may aid in phage selection and optimizing cocktail formulation based on phage life history traits. This study highlights the need for thorough characterization of phage growth parameters and informed combinations of phages, as random combinations could lead to undesirable outcomes.

## Introduction

Phage therapy, using bacteriophages to target and control bacterial infections, has emerged as a promising alternative to traditional antibiotics, especially in the face of rising antibiotic resistance. However, single-phage treatments are limited in their effectiveness against bacterial infections due to high genetic variability and rapid evolution of resistance. To address these challenges, phage cocktail, a combination of two or more phages, are employed to produce a pharmacologically diverse formulation for better efficacy and a broader spectrum in phage therapy [1, 2]. The preferable combination of phages in a cocktail should be genetically diverse, have the efficient killing of the target pathogen, adsorb to different receptors, and minimize and delay the emergence of resistant mutants in the bacterial population on the time scales of therapy [3, 4]. The outcome of phage interaction in a cocktail can lead to synergistic effects that include increased infectivity and a broader spectrum as well as antagonistic effects such as competition for the same receptor sites and limited cellular resources upon infecting a common host [5, 6]. While the concept of phage cocktails in phage therapy is well established, there remains a significant gap in our understanding of how short-term infection dynamics of component phages influence the performance of a cocktail.

The phage-mediated bacterial pathogen killing in phage therapy can be passive or demand phage replication in active treatment as in the case of bacteria inhabiting biofilms [7, 8]. The principal advantage of using a phage as a drug is auto-dosing and phage replication is a crucial factor that determines the efficacy of phage therapy in active treatment [9]. Thus, designing a phage cocktail with optimal therapeutic potential requires the study of the infection profile of each phage individually and in combination. Phage-phage interactions are primarily governed by the life history traits of the phages, including their adsorption rate, latency period, and burst size. Phages with the shortest latent period, large burst size, and highest adsorption rate are thought to exert a selective advantage in cocktails [10]. This can lead to a reduced opportunity for the slower phage to exert desired productive infection against the target pathogen. Additionally, an excluded phage can influence the replication cycle of other phages by reducing their yield [11]. Interestingly, traits like lysis time and burst size depend on host phenotype and can vary with growth conditions [12, 13]. Hence, studying how different growth media impact phage coinfection outcomes could offer valuable insights.

Understanding the influence of different life history traits of phages can be enhanced by studying phage population dynamics using mathematical models in conjunction with experimental data. Mathematical models on phage therapy, using *in vivo* data [14, 15] have unveiled key phage-immune synergy parameters. However, *in silico* understanding of phage coinfection dynamics in bacterial hosts remains limited. A model developed recently reported the kinetics of one bacterium and two phages with a focus on phage resistance development in the case *Pseudomonas aeruginosa* [16]. Hence, there is a scope for theoretical modeling of phage coinfection as it can supplement the experiments to provide a insights into phage coinfection.

This study was aimed at investigating the population dynamics of *E. coli* host cell, infected with different combinations of three phages, namely, N4, KKE5P, and Ec_YwIITB1, and monitored for their ability to efficiently reduce the pathogen population and also delay the emergence of resistance. Furthermore, experiments were guided by a mathematical model, developed based on phage growth parameters from experimental data. Our findings show that phage N4 exhibited robust infection characteristics, inhibiting the growth of Ec_YwIITB1 in coinfection experiments and time-delay experiments, where phage N4 was introduced later. Despite its wide host range, Ec_YwIITB1 did not prove to be a suitable candidate for phage cocktail formulation due to its inability to replicate in coinfection experiments. Furthermore, the metabolic state of bacterial cells influenced the life history traits of the phages in monoinfection profiles as expected. Phage N4 remained dominant in coinfection experiments, regardless of the media composition, suggesting resource is not a limiting factor. In contrast, in N4 and KKE5P coinfections, both phages replicated and coexisted, indicating positive interaction for phage cocktail development.

Based on our findings, we propose that genetically diverse phages with similar latency periods are ideal candidates for inclusion in phage cocktails. In addition, our modeling approach complemented experimental findings and provided insights into the coinfection dynamics. This integrated approach can aid in the rational design of phage cocktails. Overall, this study highlights the importance of phage characterization and the selection of phages with compatible traits.

## Materials and Methods

### Bacteriophages, whole genome sequencing, bacterial strains, and culture conditions

*Escherichia coli* Phage N4 used in this study is a laboratory strain (NCBI Genbank accession number: NC_008720.1) supplied by Prof. Ian Molineux (University of Texas) and host *E. coli* W3350 was obtained from the Yale E. coli Genetic Stock Center (CGSC). *E. coli* Phages Ec_YwIITB1 and KKE5P were isolated in the lab against MTCC *E. coli* strains, MG1687 and W3350, respectively. Phages were purified by the polyethylene glycol (PEG) precipitation method [17], followed by genome extraction using the Norgen phage DNA isolation kit (Product # 46800). The whole genomes of both phages were sequenced using the NovaSeq 6000 Sequencing System (Illumina) and assembled using the processed Illumina data on SPAdes assembler [18]. The gene/protein prediction analysis was performed using the assembled genomes in Prokka tool [19]. Sequence data of Ec_YwIITB1 and KKE5P have been uploaded to the NCBI GenBank under the accession number PQ356294 and PQ356293, respectively. In order to determine the genomic relatedness among all phage genomes, EasyFig was used for multiple genome alignment (Figure S1) [20]. Phage receptors were predicted on the basis of their known receptors of homologs. Bacterial strains were stored at -80℃ in 25% glycerol and cultivated at 37℃ with shaking at 150 rpm in Luria Broth. Phages were propagated by infecting the overnight-grown bacterial host using the double layer agar method. Phage titers were determined by plaque assays. Phages were individually mixed with the common host bacteria (*E. coli* W3350) and incubated at 37°C for 10 min to allow adsorption of phage to the host cell. The suspension was then mixed with soft agar (0.4 % agar) and poured onto Luria Agar plates. Plates were incubated at 37°C and plaques were enumerated to determine phage titer.

### Determination of phage growth parameters Latent period and burst size

Experiments on phage growth kinetics to determine latency period and burst size of phages N4, Ec_YwIITB1, and KKE5P were performed by classical one-step growth curve analysis as per the method of Ellis & Delbrück (1939) [21] with some modifications (Figure S2 A-C) [22]. Briefly, bacterial culture at the early exponential phase was infected with phage at a multiplicity of infection (MOI) of 0.01 in a total volume of 1 ml and incubated at 37℃ for 10 min. The bacteria-phage mix was centrifuged at 6000 g and the supernatant containing free phages was discarded and the pellet was resuspended in 1 ml of LB (repeated twice). Following this, 100 µl of the resuspension was diluted in 9.9 ml of LB followed by further serial dilution into another 9.9 ml LB to prevent any new infection. Samples were taken every 10 min and plaque assays were performed to enumerate the number of phage particles. The latent period is defined as the time interval between phage adsorption to host cell and its lysis. The ratio of average pfu/ml at plateau and latent period is used to compute burst size.

### Adsorption rate

Adsorption rates of phages were determined by mixing the phages at an MOI of 0.01, followed by measuring the free phage particles in the supernatant as a function of time. N4, Ec_YwIITB1, and KKE5P were mixed individually with 1 ml of the overnight grown *E. coli* W3350 (1.1 × 10⁹ cfu/ml) at an MOI 0.01 and incubated at 37°C for 20 min. At regular intervals, samples were withdrawn and centrifuged to remove the bacteria along with adsorbed phage. The unadsorbed phages in the supernatant were enumerated by plaque assay. The phage adsorption rate depends on the host and the adsorption conditions. It is expressed as an adsorption constant (k) and represented using units ml/min. The adsorption rate was evaluated by linear regression line fitting of the logarithmically transformed relative phage concentration versus time (Figure S3, A-C). By assuming a constant host density during the analysis, the adsorption rate was calculated by dividing the slope value by bacterial cell density [23].

Adsorption rate was estimated by utilizing following equation: -

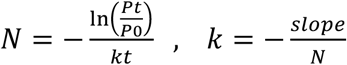

here, *Pt* - Phage titer at time t

*P0* - Phage titer of Control

*N* - Bacterial cell density

*k* - Adsorption constant to be calculated.

### Growth dynamics of the *E. coli* W3350 strain monoinfected and coinfected with N4 and Ec_YwIITB1

The experiments were conducted using transparent flat bottom 96 well plates as follows. The overnight culture of the *E. coli* W3350 strain was adjusted to OD₆₀₀ to 0.3 (which corresponded to ∼ 5×10⁷ cfu/ml) by diluting it in the prewarmed LB media. The N4 and Ec_YwIITB1 phage lysate concentrations were adjusted to the desired phage titer by diluting in LB media per the experimental conditions. For each condition, 100 µl (10^6^ cfu/ml) of the adjusted bacterial inoculum was infected with phage N, Ec_YwIITB1, and a combination of both to achieve an MOI of one in a total volume of 200 µl per well. The bacterial inoculum without phage was used in addition to the media blank as controls. The suspension was incubated at 37°C for 10 min. The unadsorbed phages were removed by centrifugation, and the pellet was resuspended in prewarmed LB. The suspension for each condition (N4 and Ec_YwIITB1 monoinfections and their coinfection) was placed in a 96-well microtiter plate and incubated at 37 °C with shaking in a plate reader (Multiskan, ThermoFisher). The growth was monitored by measuring OD₆₀₀ at 5 min intervals for up to 4 h. Data were recorded using SkanIt software version 6.0.2, with precision mode. At the end of the infection cycle, the content of each well (each condition) was collected and treated with chloroform to lyse the cells, followed by centrifugation to remove bacterial debris. The lysates obtained were stored at 4℃ and used for further plaque assays and qPCR analysis. The growth curve under each condition was obtained by plotting the OD₆₀₀ value versus time. All assays were performed in triplicates. Similarly, N4 and Ec_YwIITB1 growth dynamics were monitored under different experimental conditions (different MOI, different ratios of N4 and Ec_YwIITB1, and various time points).

### Determination of phage DNA copy number by qPCR

Primers were designed for N4 gene *68*, Ec_YwIITB1 gene *22* and KKE5P gene *10* which encodes for large terminase (gp 68), small outer capsid protein (gp 22) and primase (gp 10) respectively (Table S1). NCBI BLAST and PrimerQuest tools were used to analyze *in silico* primer specificity. The specific amplification condition of each amplicon was optimized with multiple combinations of primer with and without template. The standard curve of the Ct value versus copy number was generated with standardized plasmids carrying the gene *68* of N4, gene *22* of Ec_YwIITB1, and gene *10* of KKE5P, all cloned into the pET28a plasmid (Figure S4). Phage lysates collected from growth curve dynamics under each condition were used as a template to determine the copy number of *g*68*, g22* and g*10*. For absolute quantification of gene copy number in each sample, 1 µl of template (plasmid carrying the gene of interest) was used in a 20 µl of reaction mixture with 250 nM of primer and 10 µl of SyBr Green Master Mix (PowerUp™ SYBR™ Applied Biosystems). The qPCR cycling condition used was as follows: one cycle at 50 ℃ for 2 min, one cycle at 95℃ for 10 min and 30 cycles of (95 °C for 30 s, 62 °C for 1 min) followed by melt curve analysis (one cycle of 95 °C for 15 s, 60 °C for 1 min, and 95 °C for 15 s). Ct values for each experimental conditions were determined and used for absolute quantification of gene copy number using the standard curves (Figure S4 A-C).

### Phage infection dynamics in different media

The growth dynamics of the *E. coli* W3350 strain monoinfected and coinfected with N4 and Ec_YwIITB1 were examined in different types of bacterial culture media, namely, M9 minimal media, Luria broth, terrific broth, and super broth (their compositions are given in Table S2). For determining the growth rate of uninfected *E. coli* cells in growth media, 1% overnight grown culture was inoculated in each well consisting of 200 µl of each type of media in triplicates. Plates were incubated at 37°C with shaking in a plate reader (Multiskan, ThermoFisher), and growth was monitored by measuring OD₆₀₀ at 15-minute intervals for 16 h. The growth rate was determined by fitting the growth curve using the Slogistic1 growth model in OriginPro 2021, a data analysis and graphing software (Table S3). Similarly, mono and coinfections of host cells in different media were performed in transparent flat bottom 96- well plates at MOI 1. Phage lysates corresponding to each media condition were analyzed through qPCR and plaque assay to determine phage titers. *E.coli* cell size was determined in different media by fixing cells onto a coverslip with 2.5% glutaraldehyde, followed by dehydration by graded ethanol and drying with hexamethyldisilazane (HMDS). The coverslip was mounted on a metal stub using a sticky carbon disc and sputter coated with platinum for 2 minutes followed by imaging using a Field-emission-gun-based scanning electron microscope (JSM 7600F) (Figure S6).

### Infection dynamics of single phage and phage cocktail to determine resistance evolution

Growth dynamics of the *E. coli* W3350 strain monoinfected and coinfected with N4, Ec_YwIITB1, and KKE5P were examined for 24 h. *E. coli* cells (10^7^ cfu/ml) were monoinfected with N4, Ec_YwIITB1, KKE5P, and coinfected with N4+Ec_YwIITB1 and N4+KKE5P at MOI 1 (Figure S7). The plates were incubated at 37°C with shaking in a plate reader (Multiskan, ThermoFisher), and growth was monitored by measuring OD₆₀₀ at 20 min intervals for 24 h. The proportion of replicates that remain sensitive and show regrowth was estimated for each infection condition. The Principal Component Analysis (PCA) was performed on OD600 trajectories to determine the value of PC1, which was used to score the emergence of resistance (Figure S8). Phage-resistant mutant strains were isolated from the phage by multiple rounds of streaking. A 1% of the overnight-grown culture of resistant strains was inoculated in each well consisting of 200 µl of LB along with wild-type, and the growth rates were determined. The growth rates of resistant strains were determined by fitting the growth curve using the Slogistic1 growth model in the OriginPro 2021 software. Fitness cost, imposed by resistance mutations under various phage treatment, was accessed relative to ancestral bacterial strains by comparing the growth of isolated resistant mutants to ancestral *E.coli* host. Relative fitness was calculated as a ratio of the growth rate of resistant mutant to ancestor bacteria, where 1 indicates fitness equivalent to ancestral bacteria while fitness < 1 indicates reduced fitness.

### Monoinfection model

The monoinfection model includes three equations (Equations, 1-3) describing the dynamics of the susceptible host cells (*T*), virus (*V*), and infected cells (*I*). Susceptible cells grow logistically 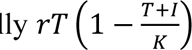 with a rate *r* and maximum capacity *K*. Virus infects susceptible cells at a rate *α*, while infected cells produce new virus particles at a rate *b* and die at a rate *d*.

Virus loss is accounted for by their “attachment” to host cells, and virus decay was assumed negligible due to the very short duration of the experiments.

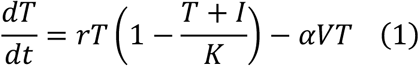

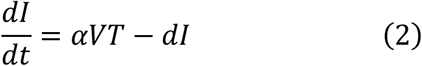

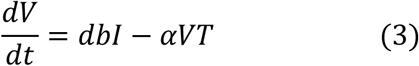

### Model non-dimensionalization

Rescaling of variables as: *T*^∗^ = *T*/*K*, *I*^∗^ = *I*/*K* , *V*^∗^ = *V*/*K* was done for both the models which simply corresponds to setting *K* = 1. As population densities are rescaled all parameters either have units of 1/time or are dimensionless. In terms of these variables, equations (1-3) become:

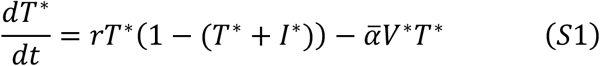

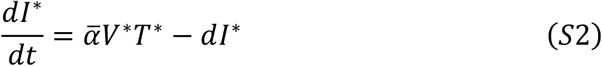

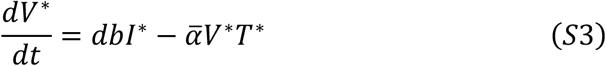

where *α*; = *αK*. All modeling plots were generated using ggplot function in R.

After conducting numerical simulations, we used a scaling factor (*K*) to convert normalized population values (*T*^∗^, *I*^∗^, *V*^∗^) to microbial concentrations (*T*, *I*, *V*). We employed a conversion formula *y* = (1.41819 × 10^9^) *x* + (-9.51032 × 10^7^) derived from the standard curve. This formula allowed us to estimate PFU/mL from the OD600 measurements, yielding a *K* value of 1.2 × 10^9^ for OD600 values (Figure S9).

### Coinfection model

The key assumptions of the coinfection model are: (1) a single host cell can only be infected by one virus, (2) no consideration of bacterial host defenses or immune responses, (3) no account for mutations, and their effects on phage-bacteria coevolution, and (4) no incorporation of bacterial resistance to phage infection due to the short experiment duration. Coinfection model of two viruses is as follows: -

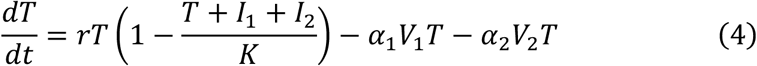

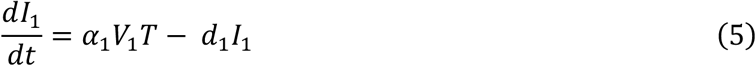

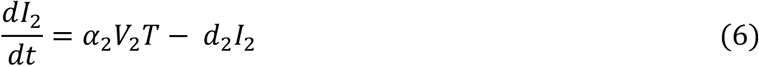

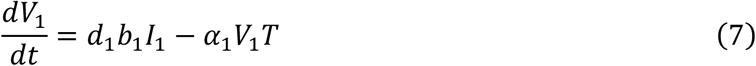

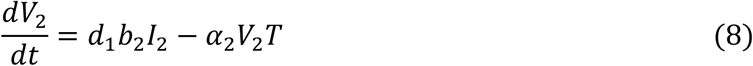

Virus *V*_2_and *V*_3_infect the susceptible target cells, *T* with rates *α*_2_and *α*_3_, respectively. The newly infected cells *I*_2_ and *I*_3_produce viruses at rates *b*_2_ and *b*_3_, and die at rates *d*_2_ and *d*_3_. Parameters estimated from monoinfection experiments were used to test the coinfection model’s ability to reproduce experimental results. The ODEs were solved using the deSolve package using R programming [24].

### Statistical significance

Statistical significance was analyzed using OriginPro 2021. The normality of the data was first checked using the Kolmogorov-Smirnov (KS) normality test. Student’s t test was used to compare means of two parametric datasets. One-way ANOVA tests were performed for parametric datasets with multiple conditions, followed by Tukey’s post-hoc test for multiple pairwise comparison. Significance is indicated in the figures by asterisks (ns, non-significant *p*-value>0.05; \**p*-value ≤ 0.05; \*\**p*-value ≤ 0.01; \*\*\**p*-value ≤0.001). Plots for experimental data and modeling were curated using OriginPro 2021 and R, respectively.

## Results

### Characterization of N4 and Ec_YwIITB1 phages

This study compared the infection outcomes of the common *E.coli* W3350 host by two unrelated lytic bacteriophages, N4 and Ec_YwIITB1. Phage N4 belongs to the *Podoviridae* family, while Ec_YwIITB1, isolated from the Yamuna River (India), belongs to the *Myoviridae* family (Figure 1A and 1B). N4 forms large and clear plaques, while Ec_YwIITB1 forms small and clear plaques on *E. coli* W3350 (Figure 1C and 1D). Life history traits such as burst size, latent period, adsorption rate, and decay rate of these phages are distinct (Figures S2 and S3; Table S4).

**Figure 1.**
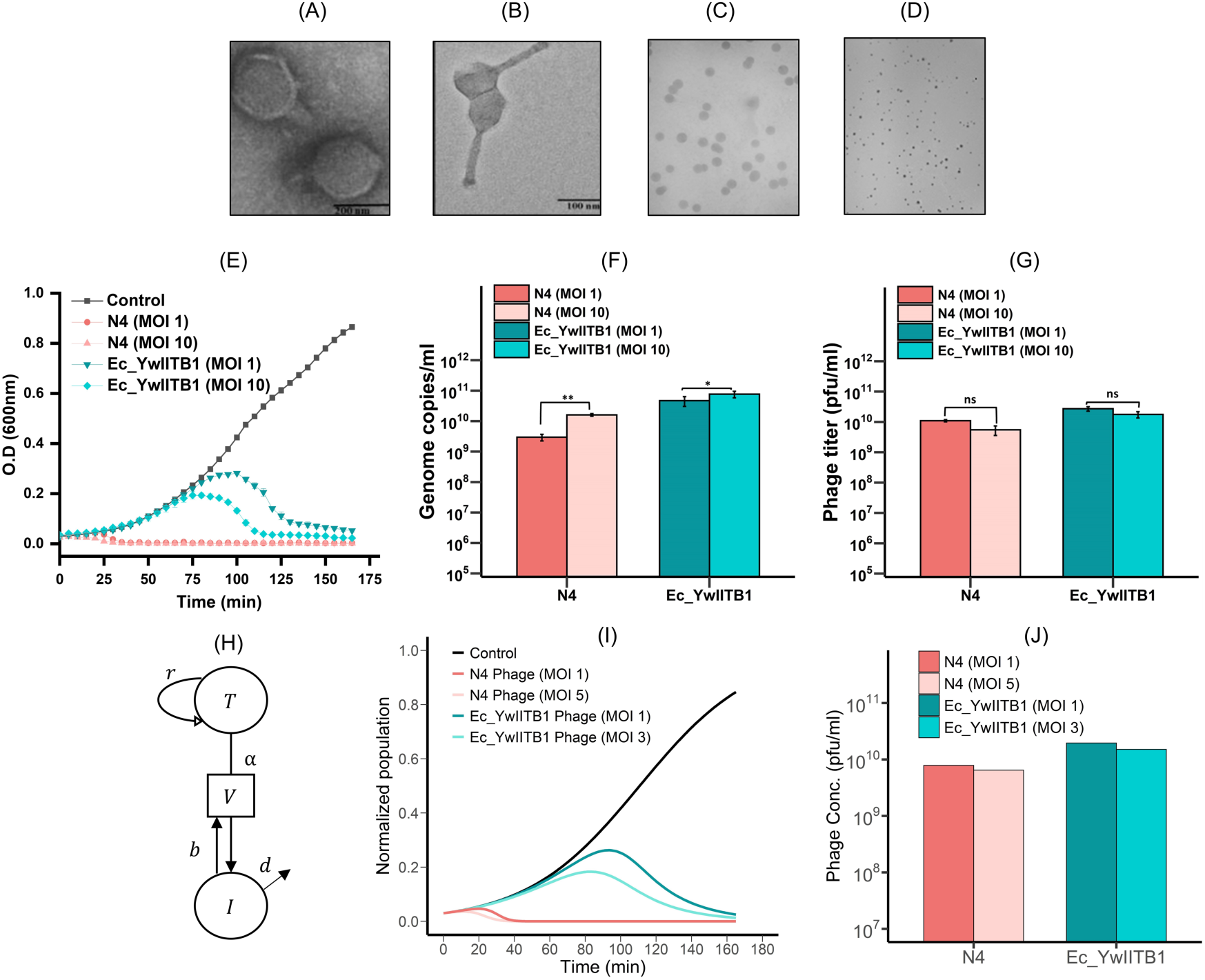
Comparative analysis of N4 and Ec_YwIITB1 phages: morphology, monoinfection dynamics, and model simulation. Transmission Electron Microscopy (TEM) micrographs of (A) N4 phage, member of the *Podoviridae* family, and (B) Ec_YwIITB1 phage that belongs to the *Myoviridae* family. Plaque morphologies of phages (C) N4 and (D) Ec_YwIITB1. (E) Optical density as a function of time of incubation. *E. coli* W3350 without phage (control), with N4 (Monoinfection), Ec_YwIITB1 (Monoinfection) at MOI of 1 and 10. Representative bar plot of (F) genome copy number estimated by absolute qPCR (G) phage titer estimated by double agar overlay method for N4 and Ec_YwIITB1 at MOI 1 and MOI 10 after one infection cycle. Statistical analysis was assessed using two-tailed student t-test for comparing means between N4 MOI 1 and MOI 10 as well as Ec_YwIITB1 MOI 1 and MOI 10 (ns, non-significant *p*-value>0.05; \**p*-value ≤ 0.05; \*\**p*-value ≤0.01) (H) Schematic diagram for the monoinfection model. In the presence of virus [V], target cells [T] which are uninfected bacterial cells get infected and moved to infected cells [I]. New viruses are released when infected cells are lysed. Capital letters represent variables and small letters indicate model parameters. (I) Phage monoinfection model simulations with the following parameters for MOI 1 simulations: ***T***_0_= 0.03, ***I***_0_= 0, ***V***_0_ = 0.03, ***r*** = 0.0314. N4 phage parameters: ***α*** = 0.0603, ***d*** = 0.5001, ***b*** = 100. Ec_YwIITB1 phage parameters: ***α*** = 0.0252, ***d*** = 0.0396, ***b*** = 50 (Table S5). Parameters for experimental MOI 10 equivalent, N4 phage: ***V***_0_ = 0.15, Ec_YwIITB1 phage: ***V***_0_= 0.09. The Y-axis depicts the sum of (***T*** + ***I***). (J) The bar plot displays phage concentration ***V*** at the end of the model simulation, scaled by ***K*** (maximum carrying capacity) with ***K*** set at 1.2 × 10^9^ pfu/ml.

### Growth curve dynamics of the *E. coli* W3350, monoinfected with N4 and Ec_YwIITB1 phages

Outcomes of phage infection of *E.coli* W3350 host were assessed by following the growth dynamics of the infected host bacterial population examined as OD₆₀₀ versus time under different conditions. At MOI 1, the growth curve of N4 and Ec_YwIITB1 infected bacterial populations displayed lysis at about 30 and 100 min, respectively, while infection at MOI 10 shortened the lysis onset time (Figure 1E). A shorter lysis time of the N4-infected cells also suggests its high virulence compared to the phage Ec_YwIITB1. This observation is consistent with the parameters extracted from the bacterial growth curves (Table S5). Lower tMax, tExt, nMax, and AUC values indicate shorter lysis times, faster attachment rates, and larger burst sizes, respectively [25]. Analysis of these variables indicates higher fitness of the N4 phage compared to the Ec_YwIITB1 against *E. coli* W3350 (Table S6). Both N4 and Ec_YwIITB1 mono infections at MOI 1 resulted in a 3-log increase in the phage titer after one infection cycle as determined by quantitative PCR (qPCR) and plaque assays. The higher burst size of N4 and delayed lysis of Ec_YwIITB1-infected cells might contribute to their high titers. However, phage titer at higher MOI doesn’t increase significantly, probably because of the abortive infection (Figure 1F and 1G).

We utilized an ODE model (Figure 1H, equations 1-3 – Materials and Methods) previously employed for the phage-bacteria system [26–29] to analyze the monoinfection dynamics based on the experimental data and predict infection outcomes under diverse conditions. After non-dimensionalization (Materials and Methods) and parameter estimation (Table S4, Figure S5), the model successfully replicated the lysis trend and phage titer at MOI 1 for both phages (Figure 1I and 1J). For MOI 10, we adjusted the initial MOI, as it reflects the effective MOI, which is lower than the actual MOI due to phage interference at entry into the host cell at higher MOIs [30]. Hence, we considered the effective MOIs of 3 and 5, respectively, for phages N4 and Ec_YwIITB1. This adjustment enabled the model to align with experimental trends for MOI 10, enhancing the predictability (Figure 1I and 1J).

### Phage coinfection dynamics with the simultaneous introduction of phages

We expanded the monoinfection model to capture the coinfection dynamics of both the phages against *E.coli* W3350 (Figure 2A, equations 4-8 – Materials and Methods). Numerical simulations of the coinfection model at MOI 1 for both the phages showed the phage N4 monoinfection trend in the growth curve (Figure 2B) and a much-reduced titer for phage Ec_YwIITB1 (Figure 2C). Increasing the MOI of phage Ec_YwIITB1 in coinfection scenarios did not substantially benefit the phage, and phage N4 remained dominant under these conditions.

**Figure 2.**
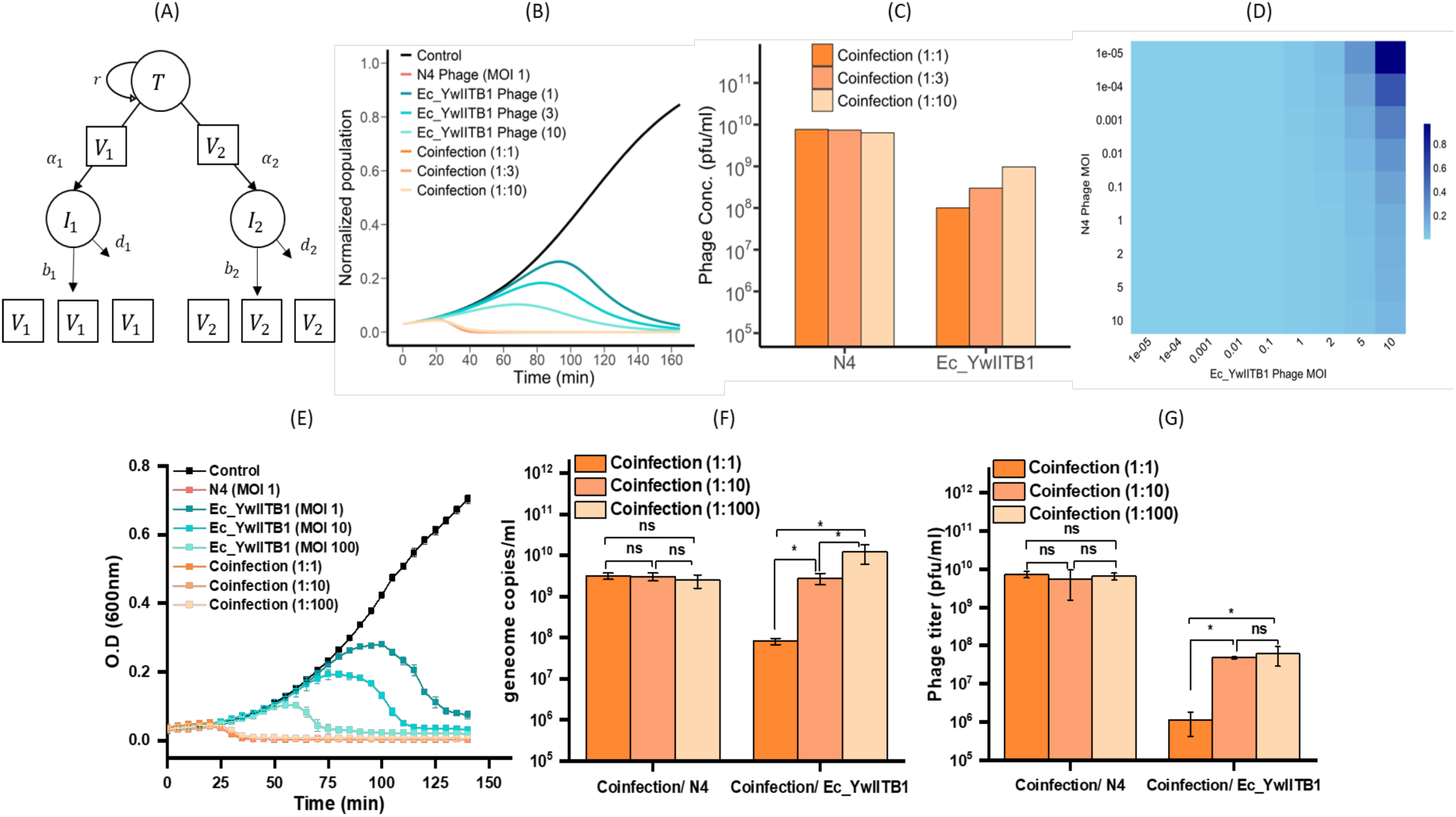
Coinfection dynamics of N4 and Ec_YwIITB1 phages: model simulations and experimental analysis. (A) The schematic diagram for the coinfection model with one host. (B) Coinfection growth curves generated from a model including uninfected host cells (control), monoinfection of N4 phage with 1 MOI, monoinfection of Ec_YwIITB1 phage with 1, 3 (equivalent to 10 MOI of experiments), and 10 (equivalent to 100 MOI of experiment) MOI. (C) Bar plot depicting phage concentrations of N4 phage and Ec_YwIITB1 phage at the end of the coinfection model simulation. (D) Heatmap illustrating the relative change in phage concentration of Ec_YwIITB1 to N4 phage (VEc_YwIITB1/VN4) when coinfected with various combinations of MOI. (E) Analysis of population dynamics of *E. coli* W3350 without phage (control), with N4 (Monoinfection), Ec_YwIITB1 (Monoinfection) at MOI 1, 10 and 100 and coinfection with different ratios of N4: Ec_YwIITB1 (1:1, 1:10, 1:100). Representative bar plot of (F) gene copy number estimated by absolute qPCR (G) phage titer estimated by double agar overlay method in case of coinfection with different ratio of Ec_YwIITB1 with respect to N4. Statistical significance was assessed between various coinfection conditions for N4 and Ec_YwIITB1 by using one-way ANOVA followed by Tukey’s test for multiple pairwise comparisons (ns, non-significant *p*-value>0.05; \**p*-value ≤ 0.05).

Varying MOI in coinfection simulations revealed that efficient replication of phage Ec_YwIITB1 during coinfection requires a substantially higher MOI of phage Ec_YwIITB1 along with a significantly lower MOI of phage N4 (for example, an Ec_YwIITB1 to N4 MOI ratio of 10: 10^-5^, Figure 2D). However, such low MOIs may not be relevant for phage therapy, as it will postpone bacterial lysis and allow bacteria to proliferate. This insight from the model suggests that the MOI of both phages needs to be optimized to enable replication of both phages without delaying the lysis of either phage.

We conducted coinfection experiments to validate the model predictions and explore the coinfection dynamics of N4 and Ec_YwIITB1 phages. Consistent with the modeling outcome, coinfection of N4 and Ec_YwIITB1 at MOI 1 followed the N4 monoinfection trend, and even at higher MOIs of Ec_YwIITB1 to N4 of 1:10 and 1:100, the lysis trend was similar to that of the N4 monoinfection (Figure 2F). While the titer of N4 was comparable to monoinfection even during coinfection (Figure 1F and Figure 2G, p-value > 0.5; not significant), that of Ec_YwIITB1 was reduced by about three-fold, suggesting a clear advantage for phage N4 (Figure 1F and Figure 2G, p-value ≤ 0.01). Correspondingly, the presence of Ec_YcIITB1 phage at a 100-fold higher MOI during coinfection did not affect the yield of N4 phage. Also, an increase in the MOI of Ec_YwIITB1 phage did not significantly increase its titer compared to its monoinfection (Figure 2G and 2H). The delay in lysis when infected with Ec_YwIITB1 appears to allow bacteria to grow for some time, while during N4 phage infection, no significant increase in the cell number before the lysis was observed. As both phages bind to the host cell using different receptors on the cell surface with similar adsorption rates (Figure S2), competition for adsorption doesn’t hold. Thus, early lysis of infected cells after completion of N4 replication and a subsequent increment in numbers (Figure 2F and 2G) seems to be the possible reason for its dominance.

### Phage coinfection dynamics with sequential addition of phages

Further, to allow Ec_YwIITB1 phage to replicate and amplify, the delayed addition of N4 was explored by modeling and experimentation. We introduced N4 phage at different time points post Ec_YwIITB1 phage infection to understand whether delayed addition could favor Ec_YwIITB1 phage propagation. First, we ran the sequential coinfection simulations using the events function from the deSolve package. Only when the addition of phage N4 was delayed by 70 min resulted in an increased titer of Ec_YwIITB1 phage (Figure 3A and 3B). Consistent with the model prediction, the delayed addition of phage N4 for up to 30 min followed the N4 monoinfection lysis trend (Figure 3C). The phage titer for Ec_YwIITB1 at these time points showed a gradual increase with increasing time delay (Figure 3E). A 70-minute delay in N4 phage introduction allowed phage Ec_YwIITB1 to complete its infection cycle, leading to a 3- log increase in its titer and much reduced N4 phage titer (Figure 3D and 3E). Since the sequential addition of phages is not a common phage therapy practice, our novel insight highlights the importance of selecting phages with thorough characterization of life history traits, emphasizing the importance of phages having comparable latency periods and lysis times.

**Figure 3.**
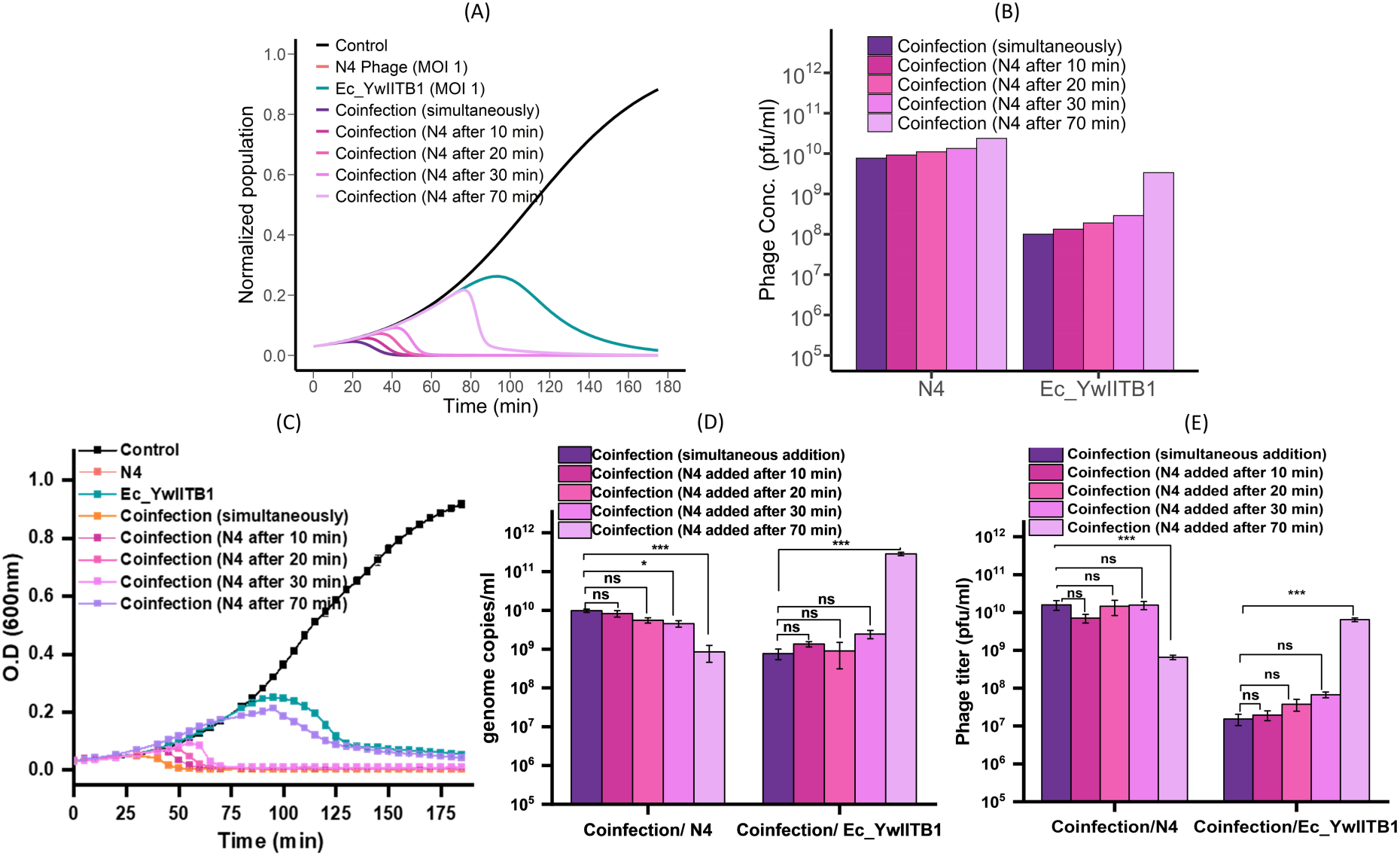
Phage coinfection dynamics with sequential introduction of phages. (A) Computational growth curve depicting the delayed introduction of N4 phage (MOI 1) in the coinfection model (Ec_YwIITB1 phage MOI 1). (B) Bar plot illustrating phage concentrations from the coinfection model with delays of 10, 20, 30, and 70 minutes in the introduction of N4 phage. (C) Analysis of population dynamics of *E. coli* W3350 without phage (control), with delayed introduction of N4 phage at different time points (10, 20, 30, and 70 min) after Ec_YwIITB1. Representative bar plot of (D) gene copy number estimated by absolute qPCR (E) phage titer estimated by double agar overlay method with delay in N4 phage addition after Ec_YwIITB1. Statistical significance was assessed by using one-way ANOVA followed by Tukey’s test for multiple pairwise comparison (ns, non-significant *p*-value>0.05; \**p*-value ≤ 0.05; \*\*\**p*-value ≤0.001).

### Phage coinfection dynamics with phages of a similar latent period

Our modeling and experimental results have shown that two phages with different latency periods can coexist only under certain conditions and may not be ideal for cocktail preparation. Hence, we explored one of the other isolates from our collection, phage KKE5P, which has a latency period of 20 min (similar to phage N4) and a burst size of about 200 on *E.coli* W3350 (Figure S1C). The adsorption rate of KKE5P was estimated to be 1.21 x 10^-10^ ml/min (Supplemental Figure S2C). Phage KKE5P possesses a capsid of 40 nm in diameter encasing a 39-kb double-stranded DNA genome and belongs to the *podoviridae* family (Figure 4A). The plaque morphology is also similar to that of phage N4 (Figure 4B). Based on its similarity to other phage T7, lipopolysaccharide (LPS) appears to be one of the host cell receptors for binding of KKE5P phage [31]. KKE5P phage monoinfection of *E.coli* W3350 follows the N4- like trend (Figure 4C). The coinfection with N4 also exhibits the same profile (Figure 4C). The qPCR estimates a 5-log increase in the phage titer for KKE5P monoinfection. During coinfection, both KKE5P and N4 phages exhibited productivity similar to their monoinfection (Figure 4D), indicating that both phages can replicate efficiently, unlike the N4 and Ec_YwIITB1 combination. This result suggests that because of a similar latent period and different host receptors, both phages can replicate effectively. Also, the coinfection lysis trend points towards a probable synergy between phages N4 and KKE5P (Figure 4C). The mathematical model, using parameters estimated from monoinfection data (Table S6) also showed that both phages could replicate successfully in a coinfection scenario (Figure 4C and D).

**Figure 4.**
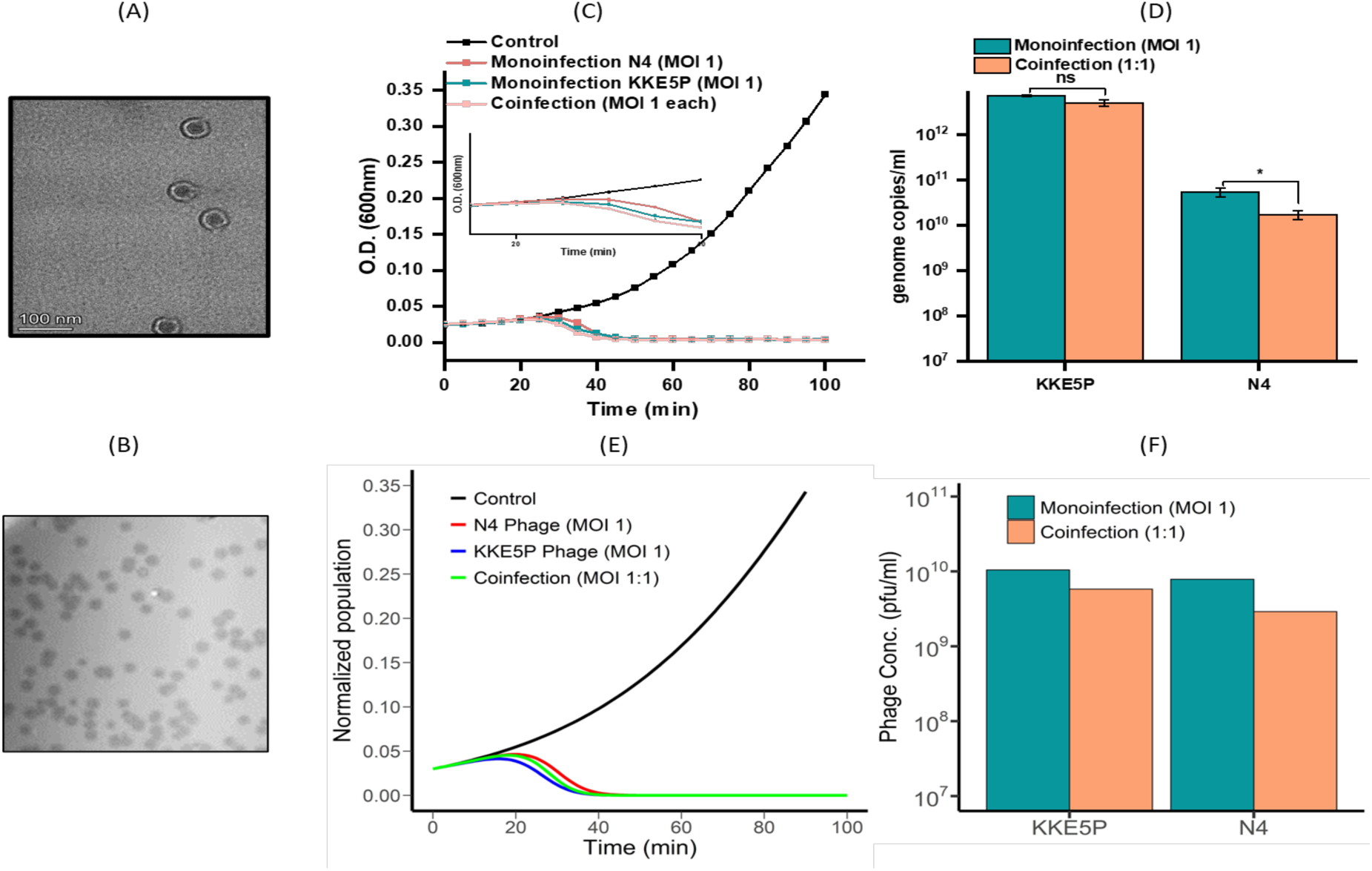
Comparative analysis of *E. coli* growth and phage infection dynamics of KKE5P and N4 with similar latent periods. (A) Transmission Electron Microscopy (TEM) micrographs of KKE5P phage belonging to *Podoviridae* family (B) Plaque morphology of KKE5P phage against *E. coli* W3350 on Luria Agar plates with O.4% top agar (C) Analysis of population dynamics of *E. coli* W3350 without phage (control), with KKE5P (Monoinfection), N4 (Monoinfection) at MOI 1 and coinfection at 1:1 ratio of KKE5P: N4. (D) Representative bar plot of gene copy number estimated by absolute qPCR for KKE5P and N4 at MOI 1 in case of monoinfection and coinfection after one infection cycle. Statistical analysis was assessed using two tailed student’s t-test for comparing means between monoinfection and coinfection of KKE5P and N4 (ns, non-significant *p*-value>0.05; \**p*-value ≤ 0.05). (E) Growth curves generated from a model including uninfected host cells (control), monoinfection of N4 phage and KKE5P phage with 1 MOI, and coinfection of both phages with 1 MOI. Parameters for MOI 1 simulations: ***T***_0_= 0.03, ***I***_0_ = 0, ***V***_0_ = 0.03, ***r*** = 0.0314. N4 phage parameters: ***α*** = 0.0603, ***d*** = 0.5001, ***b*** = 100. KKE5P phage parameters: ***α*** = 0.0503, ***d*** = 0.05497, ***b*** = 140 (Table S3). (F) Bar plot depicting phage concentrations of N4 phage and KKE5P phage at the end of the monoinfection and coinfection model simulation.

### Phage infection dynamics in different culture media

We also investigated the effect of different bacterial growth media, namely, nutrient-rich terrific broth (TB), super broth (SB), and nutrient-deficient M9, on the physiological state (cell size) of the host *E.coli* strain W3350 as it could impact phage productivity. The bacterial growth rate and doubling time varied widely in different nutrient media (Table S5, Figure 5A). Mono and coinfection experiments were also conducted in these three media (Figures 5B-D). Both N4 and Ec_YwIITB1 monoinfection displayed distinct lysis times in different growth media. Even upon providing nutrient-rich media (TB and SB), coinfection of N4 and Ec_YwIITB1 at MOI 1 resulted in a lysis trend similar to that of the N4 monoinfection (Figure 5B-D), suggesting that the observed coinfection dynamics are independent of the host growth. The productivity of both phages during coinfection was much higher in SB compared to other media (Figure 5E and 5F). Host lysis time and phage burst size are dictated by the host growth rate and physiological status [12, 32]. Apart from the growth rate, a factor attributed to an increase in phage burst size is the cell size. The Scanning Electron Microscopic (SEM) analyses of host bacterium grown in different media showed that cell sizes were larger in nutrient-rich growth media (Figure 5G). Thus, a larger cell with more resources positively influences the phage titer (Figure S6).

**Figure 5.**
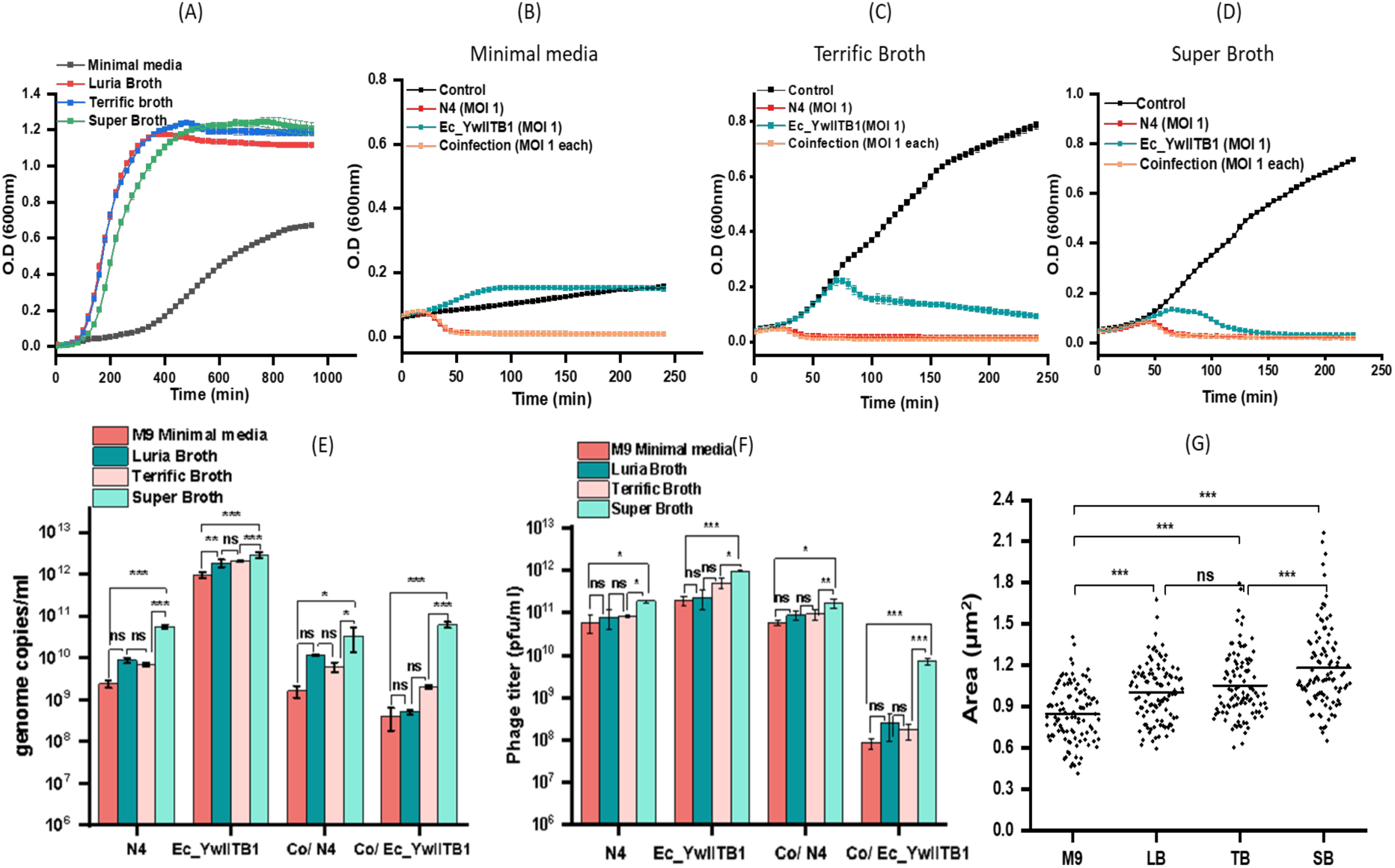
*E. coli* growth and phage infection dynamics in different culture media. (A) Growth curve of uninfected *E. coli* cells cultured in Luria-Bertani Broth, Terrific Broth, M9 Minimal media, and Super Broth cultured in 96 well plates at 37℃ for 24 hours with continuous shaking. *E. coli* growth curve dynamics under different conditions: without phage (control), with N4 phage (monoinfection), with Ec_YwIITB1 phage (monoinfection), and coinfection with N4 and Ec_YwIITB1 phage, each at MOI of 1, in (B) M9 minimal media (C) Terrific Broth (D) Super Broth. (E) Bar plot of gene copy number estimated by absolute qPCR (F) Bar plot of phage titer estimated by double agar overlay method for N4 and Ec_YwIITB1 monoinfection and coinfection at MOI 1 in different culture media. Statistical significance was assessed by using one-way ANOVA followed by Tukey’s test for multiple pairwise comparisons (ns, non-significant *p*-value>0.05; \**p*-value ≤ 0.05; \*\**p*-value ≤ 0.01; \*\*\**p*-value ≤0.001). (G) Box plot representation of analysis of the difference in *E.coli* cell size depending upon host physiological state when cultured in different bacteriological media through SEM imaging, measured through ImageJ displaying an increase in cell area with the nutrient richness of media (n=100). “Co” refers to coinfection.

Furthermore, the variations observed in the Ec_YwIITB1 monoinfection growth curves in nutrient-rich media like TB and SB (Figure 5C and 5D) compared to LB (Figure 1E) were investigated using the modeling approach. As nutrient-rich media leads to larger cell sizes (as shown in Figure 5G) that provide more resources for phage production, influencing phage burst size and infection dynamics. To reflect these conditions, we adjusted the model parameters by increasing the burst size (*b*) and decreasing the death rate (*d*) of infected cells. These adjustments aligned the simulations with the experimental observations (Figure 5C and Figure 6A– Super Broth curve). This is further supported by the phage titer data (Figure 5F, 5G, and 6B). The difference in phage productivity in nutrient-rich media has also been reported before [33].

**Figure 6.**
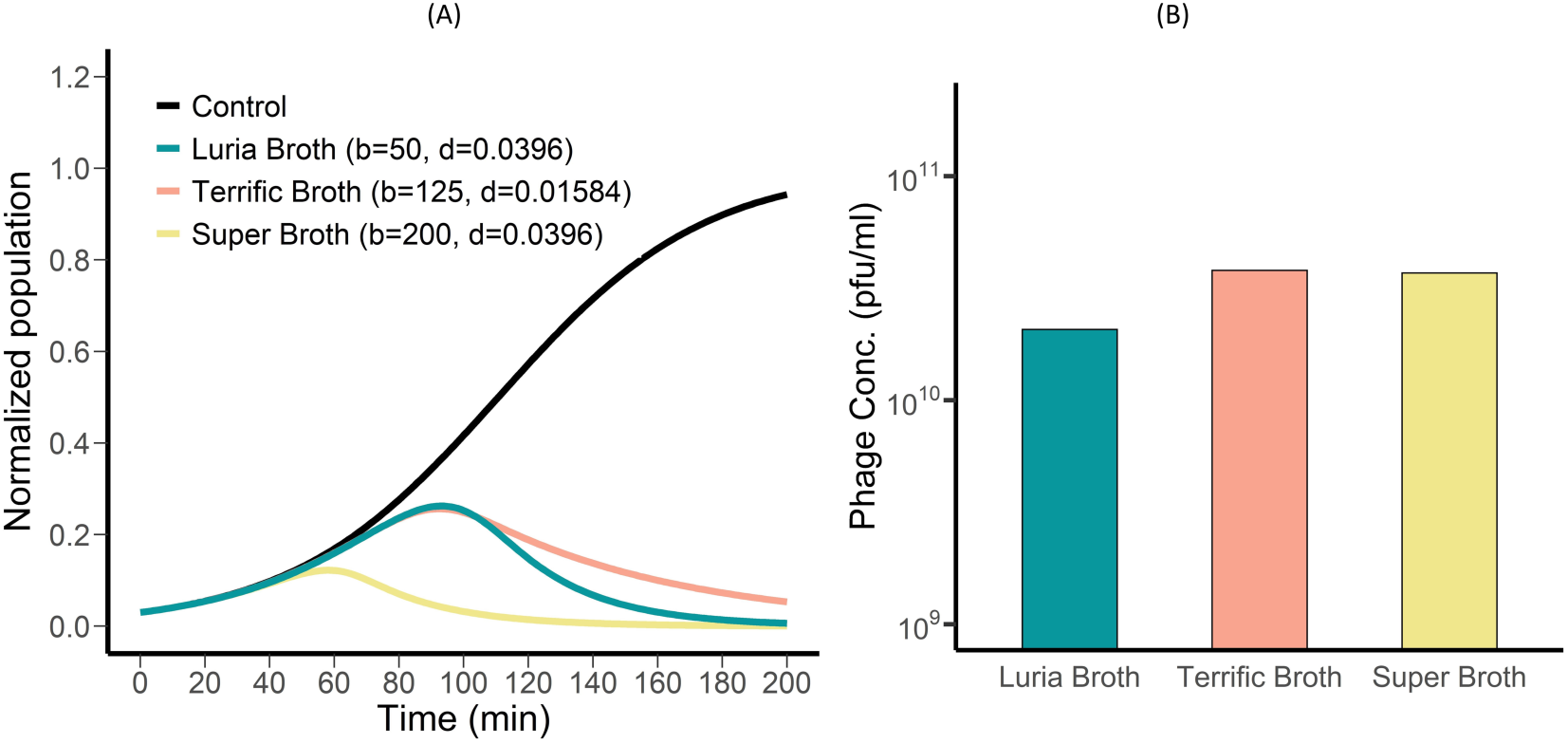
Exploration of media-induced variation in virus life history traits using a mathematical model: The control curve shows a linear increase over time and stabilizes at a normalized population value of 1. The Luria Broth curve (*b* = 50, *d* = 0.0396), corresponding to experimental results presented in Figure 1E, demonstrates a growth pattern of Ec_YwIITB1 phage at MOI1. The Terrific Broth curve (*b* = 125, *d* = 0.01584), as seen in Figure 5C, shows an extended decline, indicating the effects of increased burst size and reduced infected cell death rate. The Super Broth curve (*b* = 200, *d* = 0.0396), consistent with observations of the EcYwIITB1 growth curve from Figure 5D, reaches a lower maximum and declines earlier compared to other media conditions.

Additionally, increasing the burst size (*b*) while keeping other parameters constant led to faster curve decay (Figure 6A – Super broth curve) and a higher phage titer (Figure 6B), aligning with observations from Ec_YwIITB1 phage infection in Super Broth (Figure 5D). This suggests that larger, more metabolically active cells can influence phage life history traits and production rates. The observed variation in growth curves across different media underscores the importance of understanding phage infection profiles in various conditions before selecting them for therapeutic applications. While adjusting these parameters provided key insights into phage population dynamics, further validation of parameter identifiability is necessary to confirm their precise impact and reliability.

### Infection dynamics of single phage and phage cocktail to determine resistance evolution

The short-term experiments, conducted over approximately one infection cycle, showed the superior performance of the N4 and KKE5P combination over N4 and Ec_YwIITB1. We also assessed the coinfection dynamics of both combinations (along with monoinfection controls) for 24 h to investigate their effect on the emergence of phage resistance in infected bacterial populations. By utilizing the liquid-based assay, the difference in the OD600 values between uninfected and infected bacterial populations was used as a parameter to categorize bacteria as sensitive, partially resistant, and resistant [34]. The sensitive bacterial population displayed no regrowth, partial resistance reflected inhibited growth, while resistant bacteria showed growth similar to the uninfected bacterial population. Bacterial populations infected with a single phage, as well as in combination, were either sensitive or displayed varying degrees of partial resistance in each of the replicates (Figure S7). We characterized the proportion of sensitive and partial resistance using the data from 28 replicates for every infection condition (Figure S7). Our data show that bacterial isolates infected with individual phages N4 and KKE5P displayed almost similar partial resistant and sensitive proportions, while Ec_YwIITB1 infected bacteria showed regrowth in most replicates. Also, a significant proportion of the infected bacteria remains sensitive to the N4 and KKE5P phage combination. It highlights that the N4 and KKE5P combination is better in the suppression of resistance when compared to single phage infections confirming synergy as observed earlier (Figure 4C). In contrast, the N4 and Ec_YwIITB1 combination showed a behavior similar to N4 alone, consistent with our short-term infection dynamics (Figure 7A). To quantitatively measure resistance and to explain the variability of partial resistance, principal component analysis (PCA) was performed on the difference in the O.D trajectories of control (no phage) to monoinfected and coinfected replicates [34]. PCA reflects that PC1 accounts for 88.19% of the variance and distinguishes between sensitivity and variability of partial resistance (Figure S8). Utilizing PC1 as a measure of resistance, better suppression of resistance emergence in the N4 and KKE5P combination was found as compared to N4 or KKE5P single infection, while the N4 and Ec_YwIITB1 combination displayed a similar score as of N4 (Figure 7B). The fitness cost for the emergence of resistance in the N4 and KKE5P combination reflected much-reduced fitness. The extent of resistant phenotypes that emerged under the pressure of the N4 and Ec_YwIITB1 combination is comparable to that of the ancestral bacterial strain (Figure 7C). These findings show that the N4 and KKE5P combination is more effective in suppressing bacterial regrowth for longer periods compared to the N4 and Ec_YwIITB1 combination.

**Figure 7.**
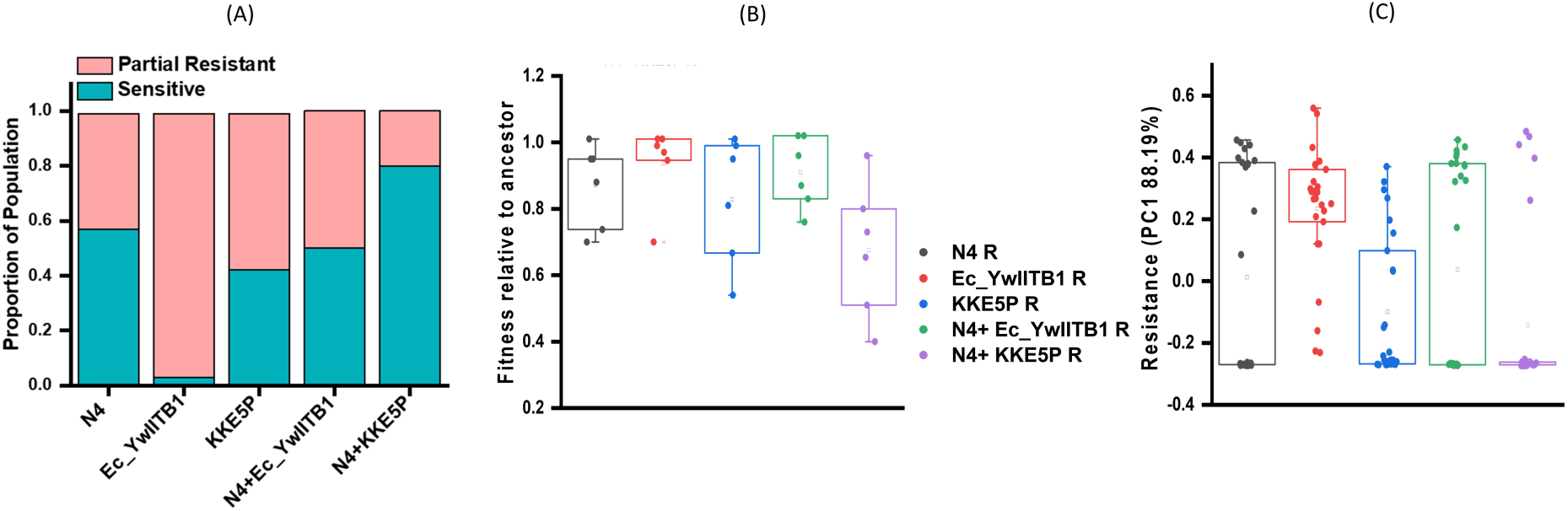
Determination of resistance evolution based on infection dynamics of single phage infection and coinfection. A) Proportion of sensitive and partial resistant bacterial isolates among 28 replicates for every infection condition displaying the least emergence of resistance for N4 and KKE5P phage cocktail. (n=27) B) Resistance measurement of various infection conditions as indicated by PC1 score of PCA reflecting bacterial isolates is less resistant for N4 and KKE5P phage cocktail. (n=27) C) Fitness of evolved resistant mutants relative to their common ancestor bacterial strain where a value of 1 indicates equivalent fitness while value below 1 indicates associated fitness costs. Resistance against N4 and KKE5P phage imposes more fitness costs relative to the ancestor in comparison to other infection conditions.

## Discussion

Phage cocktails offer a potential solution to address the limitations of single phage therapy, such as the development of resistance and polymicrobial infections [35]. Currently, there is no well-defined pipeline for the formation of effective therapeutic cocktails, although some studies have endeavored to provide a guideline for phage selection [36, 37]. The general criteria for combining phages into a cocktail is based on genomic relatedness, wide host range, different receptors, or merely, the ability to lyse bacteria. Combining genetically diverse multiple phages displays enhanced efficacy to curtail the emergence of phage resistance as phages targeting different receptors call for more fitness costs in the evolution of resistant bacteria [38, 39]. These guidelines provide limited information about the influence of the latent period on the productive infection efficacy of constituent phages in a cocktail upon coinfection. The critical parameters that define the success of phage therapy are a reduction in the bacterial count that can be handled by the immune system and delayed emergence of resistance. It is important to note that cocktails tend to behave differently than single phages because of interactions among component phages that affect phage replication, bacterial lysis, and the emergence of resistance [6]. Hence, the ability of phages in the cocktail to replicate and attain killing titers and retain prolonged infectivity are the primary determinants of efficacy.

This study explored the idea of developing a minimum-sized phage cocktail that serves the purpose of enhanced therapeutic efficacy in comparison to a single phage. The addition of too many phages can further create complexities in manufacturing, cost, and reduced dose as well as in immune response [10, 40]. The infection dynamics of phage combinations consisting of two phages (N4 and Ec_YwIITB1 and N4 and KKE5P) were studied by following the time-kill kinetics and phage replication during monoinfection and coinfection of the common host. N4 and Ec_YwIITB1 phages were combined to complement the higher virulence (lysis rate and burst size) of N4 with the broad host range of Ec_YwIITB1. Compared to monoinfection, coinfection growth curve and phage titers showed the dominance of N4 phage with no significant benefit of including the Ec_YwIITB1 phage. This suggests that the N4 and Ec_YwIITB1 combination to target bacterial resistance is ungainly because of the low-killing titer of Ec_YwIITB1. The only condition under which the coexistence of both phages is seen is the delayed introduction of N4 (Figure 3C-E) which would allow the completion of at least one infection cycle of phage Ec_YwIITB1. Since the probable reason for N4 productive infection appeared to be its shorter latent period compared to Ec_YwIITB1, we investigated the coinfection dynamics of N4 and KKE5P with a similar latent period. Although KKE5P is a podovirus with life history traits similar to phage N4, it is not closely related to phage N4 and has a different receptor for infection (Figure 4 and S10). In both monoinfection and coinfection, both phages displayed similar lysis trends and productive infection. This emphasizes the importance of using phages with a similar latency period to ensure their coexistence for the entire duration of therapy. Also, monoinfection at higher MOI led to decreased phage titers, underscoring the impact of MOI on phage production. The negative correlation observed between burst size and MOI is most likely associated with “phage-induced lysis from without” cells are overburdened and resource-constrained, resulting in inefficient phage assembly that leads to high-multiplicity-associated abortive infections and ultimately affects the phage production [41, 42].

While experimental approaches are essential for elucidating the overall efficacy and practical aspects of phage therapy, mathematical modeling offers valuable insights into the underlying mechanisms of bacteria phage infection dynamics [14, 43, 44]. A mathematical model developed in this study effectively captured bacterial growth dynamics during monoinfection for all phages, demonstrating its robustness in replicating infection patterns. Furthermore, the model offered predictive insights into coinfection outcomes, both in cases of simultaneous and sequential phage introduction, which guided the design of subsequent experiments. By varying phage MOI, the model allowed us to explore combinations that enabled Ec_YwIITB1 to coexist with N4, offering insights into coinfection dynamics that would be challenging and time-consuming to replicate experimentally. This insight from modeling was experimentally confirmed, and informed to select a phage with life history traits similar to N4 (KKE5P phage) from our collection.

Infection of a host cell by multiple phages can result in competition for limited cellular resources with adverse effects on phage production [45]. The infection dynamics of the N4- Ec_YwIITB1 combination were investigated in different bacteriological media to gain insights into how cellular resources influence the outcome of coinfection. Our results show that the lysis time and burst size of phages depend on the host metabolic status which was largely governed by the media composition [46, 47]. Bacterial cells in Super Broth displayed increased cell size and high metabolic machinery, leading to higher burst size and phage titer indicating that higher phage titers can be achieved by utilizing rich culture media like Super Broth. Furthermore, by adjusting the model parameters, we could replicate the variation in bacterial growth curves observed for phage Ec_YwIITB1 monoinfection across different media. Different types of growth media likely influenced the metabolic rate of the bacterial cells, leading to changes in key phage life history traits such as burst size and latent period. While these effects were evident in the experimental virus titers, the model provided a clearer understanding of how nutrient availability could directly impact these traits. However, it is important to note that the structural identifiability of these parameters has not been thoroughly assessed and requires further investigation. Nonetheless, the observed coinfection dynamics in LB media, the dominance of phage N4, remained the same in all types of media tested (Figure 5B-D), suggesting that the N4 dominance is because of its relatively superior infection parameters (latency period and burst size) rather than the host metabolic status or the growth conditions.

While the short-term experiments helped in gaining insights into the performance of individual phages in a cocktail, the long-term experiments (24-hour duration) using both combinations further helped in gaining insights into how phage life history traits interact and influence the emergence of resistance in bacterial populations. While the combination of N4 and KKE5P synergize and display extended suppression of resistance, the N4 and Ec_YwIITB1 behave similarly to N4 single phage as observed in the short-term experiments (Figure 2E). The slow growth rate of resistant mutants suggests a higher fitness cost imposed by the combination of N4 and KKE5P phages. This suggests that phage cocktails can delay the emergence of resistance only if constituent phages can persist in high titers during therapy. Extensive analysis of genomes of resistant strains is required to determine the type of mutations imposing the fitness burden on the phage-escape mutants. However, as observed in the experimental outcomes (Figure S7), the resistance dynamics demonstrated stochastic behavior. The ODE-based model could not capture this outcome, indicating that a deterministic approach is not fully adequate for modeling these resistance mechanisms. Furthermore, the context- dependance of the interplay between phage-phage-host interactions, mutation rates, and fitness costs associated with resistance emergence, making it challenging to identify a clear set of parameters that could consistently explain the observed outcomes. As a result, it is beyond the scope of the current model to predict the emergence or reduction of resistance in different coinfection scenarios. Future work would require incorporating stochastic modeling approaches and evolution dynamics to better account for the random nature of resistance development in bacterial populations.

In a few randomized clinical trials, patients receiving the phage cocktail showed no improvement in the clinical outcome compared to the placebo, because of the lack of achieving productive infection by phages [48, 49]. Our study, in line with this clinical observation, suggests that arbitrary combinations of phages for therapy may exhibit uncertain outcomes because of the interaction among component phages. Thus, phages must be characterized before combining to elucidate their infection kinetics for productive infection in combination with other phages to form a minimum-size cocktail. Understanding the phenomenon of phage coinfection during short-term dynamics for one group of phages from this study would upskill the knowledge of other phages. Additionally, the mathematical model can test hypotheses about different conditions a priori to experimental studies and assist in phage selection for cocktail design.

## Acknowledgments

Research in the KK lab is supported by the Department of Science and Technology, DST [Indo- RSF, DST/IC/RSF/2024/460) and Council of Scientific and Industrial Research (CSIR, 37/1752/23/EMR-II). R.K. acknowledges the financial support provided by the IIT Bombay Senior Research Fellowship. H.V.C. acknowledges financial support provided by the DST INSPIRE Senior Research Fellowship.

## CReDiT

**RK**: Conceptualization, Methodology, Investigation. Formal analysis, Writing-Original draft preparation, Writing- Review & Editing; **HVC:** Conceptualization, Methodology, Investigation, Resources, Formal analysis, Writing-Original draft preparation, Writing- Review & Editing, **MMI**: Methodology, Investigation, Resources, Formal Analysis, Writing- Editing, **KK:** Conceptualization, Methodology, Formal analysis, Writing-Original draft preparation; Writing- Review & Editing, Supervision, Funding acquisition.

## Data Availability

All data generated or analyzed during this study are included in this published article and its supplementary information files.

## References

1. Ferry T, Kolenda C, Laurent F, Leboucher G, Merabischvilli M, Djebara S, Gustave CA et al. Personalized bacteriophage therapy to treat pandrug-resistant spinal Pseudomonas aeruginosa infection. Nature communications. 2022 Jul 22;13(1):4239. 10.1038/s41467-022-31837-9

2. Federici S, Kredo-Russo S, Valdés-Mas R, Kviatcovsky D, Weinstock E, Matiuhin Y et al. Targeted suppression of human IBD-associated gut microbiota commensals by phage consortia for treatment of intestinal inflammation. Cell. 2022 Aug 4;185(16):2879–98. 10.1016/j.cell.2022.07.003

3. Tanji Y, Shimada T, Yoichi M, Miyanaga K, Hori K, Unno H. Toward rational control of Escherichia coli O157: H7 by a phage cocktail. Applied Microbiology and Biotechnology. 2004 Apr;64:270–4. 10.1007/s00253-003-1438-9

4. Merabishvili M, Pirnay JP, De Vos D. Guidelines to compose an ideal bacteriophage cocktail. Bacteriophage Therapy: From Lab to Clinical Practice. 2018:99–110. 10.1007/978-1-4939-7395-8_9

5. Schmerer M, Molineux IJ, Bull JJ. Synergy as a rationale for phage therapy using phage cocktails. PeerJ. 2014 Sep 25;2:e590. 10.7717/peerj.590

6. Niu YD, Liu H, Du H, Meng R, Sayed Mahmoud E, Wang G et al. Efficacy of individual bacteriophages does not predict efficacy of bacteriophage cocktails for control of Escherichia coli O157. Frontiers in Microbiology. 2021 Feb 24;12:616712. 10.3389/fmicb.2021.616712

7. Harper DR, Parracho HM, Walker J, Sharp R, Hughes G, Werthén M, et al. Bacteriophages and biofilms. Antibiotics. 2014 Jun 25;3(3):270-84. 10.3390/antibiotics3030270

8. Chan BK, Abedon ST. Phage therapy pharmacology: phage cocktails. InAdvances in applied microbiology 2012 Jan 1 (Vol. 78, pp. 1-23). Academic Press. 10.1016/B978-0-12-394805-2.00001-4

9. Loc-Carrillo C, Abedon ST. Pros and cons of phage therapy. Bacteriophage. 2011 Mar 1;1(2):111–4. 10.4161/bact.1.2.14590

10. Nilsson AS. Pharmacological limitations of phage therapy. Upsala journal of medical sciences. 2019 Oct 2;124(4):218–27. 10.1080/03009734.2019.1688433

11. Delbrück M. Interference between bacterial viruses: III. The mutual exclusion effect and the depressor effect. Journal of bacteriology. 1945 Aug;50(2):151–70.

12. Hadas H, Einav M, Fishov I, Zaritsky A. Bacteriophage T4 development depends on the physiology of its host Escherichia coli. Microbiology. 1997 Jan;143(1):179–85. 10.1099/00221287-143-1-179

13. Demory D, Arsenieff L, Simon N, Six C, Rigaut-Jalabert F, Marie D et al. Temperature is a key factor in Micromonas–virus interactions. The ISME journal. 2017 Mar;11(3):601–12. 10.1038/ismej.2016.160

14. Roach DR, Leung CY, Henry M, Morello E, Singh D, Di Santo JP. Synergy between the host immune system and bacteriophage is essential for successful phage therapy against an acute respiratory pathogen. Cell host & microbe. 2017 Jul 12;22(1):38–47. 10.1016/j.chom.2017.06.018

15. Delattre R, Seurat J, Haddad F, Nguyen TT, Gaborieau B, Kane R et al. Combination of in vivo phage therapy data with in silico model highlights key parameters for pneumonia treatment efficacy. Cell Reports. 2022 May 17;39(7). 10.1016/j.celrep.2022.110825

16. Yu Z, Luong T, Banuelos S, Sue A, Horn MA, Ryu H, Roach D, Segal R, Huang Q et al. Modeling multiphage-bacteria kinetics to predict phage therapy potency and longevity. bioRxiv. 2022 Nov 11:2022–11. 10.1101/2022.11.11.516137

17. Sambrook, Joseph. Molecular Cloning : a Laboratory Manual. Cold Spring Harbor, N.Y. :Cold Spring Harbor Laboratory Press, 2001

18. Andrey P, Dmitry A, Dmitry M, Alla L, Anton K. Using SPAdes de novo assembler. Curr Protoc Bioinformatics. 2020;70(1):e102. 10.1002/cpbi.102

19. Seemann T. Prokka: rapid prokaryotic genome annotation. Bioinformatics. 2014 Jul 15;30(14):2068–9. 10.1093/bioinformatics/btu153

20. Sullivan MJ, Petty NK, Beatson SA. Easyfig: a genome comparison visualizer. Bioinformatics. 2011 Apr 1;27(7):1009–10. 10.1093/bioinformatics/btr039

21. Ellis EL, Delbruck M. The growth of bacteriophage. The Journal of general physiology. 1939 Jan 20;22(3):365–84. 10.1085/jgp.22.3.365

22. Kropinski AM. Practical advice on the one-step growth curve. Bacteriophages: Methods and Protocols, Volume 3. 2018:41–7.

23. Hyman P, Abedon ST. Practical methods for determining phage growth parameters. Bacteriophages: methods and protocols, volume 1: isolation, characterization, and interactions. 2009:175-202. 10.1007/978-1-60327-164-6_18

24. Soetaert KE, Petzoldt T, Setzer RW. Solving differential equations in R: package deSolve. Journal of statistical software. 2010;33(9). 10.18637/jss.v033.i09

25. Turner PE, Draghi JA, Wilpiszeski R. High-throughput analysis of growth differences among phage strains. Journal of microbiological methods. 2012 Jan 1;88(1):117–21. 10.1016/j.mimet.2011.10.020

26. Sinha V, Goyal A, Svenningsen SL, Semsey S, Krishna S. In silico evolution of lysis-lysogeny strategies reproduces observed lysogeny propensities in temperate bacteriophages. Frontiers in microbiology. 2017 Jul 26;8:1386. 10.3389/fmicb.2017.01386

27. Bohannan BJ, Lenski RE. Linking genetic change to community evolution: insights from studies of bacteria and bacteriophage. Ecology letters. 2000 Jul;3(4):362–77. 10.1046/j.1461-0248.2000.00161.x

28. Payne RJ, Jansen VA. Understanding bacteriophage therapy as a density-dependent kinetic process. Journal of Theoretical Biology. 2001 Jan 7;208(1):37–48. 10.1006/jtbi.2000.219

29. Weitz JS, Hartman H, Levin SA. Coevolutionary arms races between bacteria and bacteriophage. Proceedings of the National Academy of Sciences. 2005 Jul 5;102(27):9535–40. 10.1073/pnas.0504062102

30. Nguyen TV, Wu Y, Yao T, Trinh JT, Zeng L, Chemla YR, et al. Coinfecting phages impede each other’s entry into the cell. Current Biology. 2024 Jul 8;34(13):2841–53. 10.1016/j.cub.2024.05.032

31. Chen W, Xiao H, Wang L, Wang X, Tan Z, Han Z et al. Structural changes in bacteriophage T7 upon receptor-induced genome ejection. Proceedings of the National Academy of Sciences. 2021 Sep 14;118(37):e2102003118. 10.1073/pnas.2102003118

32. Golec P, Karczewska-Golec J, Łoś M, Węgrzyn G. Bacteriophage T4 can produce progeny virions in extremely slowly growing Escherichia coli host: comparison of a mathematical model with the experimental data. FEMS microbiology letters. 2014 Feb 1;351(2):156–61. 10.1111/1574-6968.12372

33. Warner CM, Barker N, Lee SW, Perkins EJ. M13 bacteriophage production for large-scale applications. Bioprocess and biosystems engineering. 2014 Oct;37:2067–72. 10.1007/s00449-014-1184-7

34. Borin JM, Avrani S, Barrick JE, Petrie KL, Meyer JR. Coevolutionary phage training leads to greater bacterial suppression and delays the evolution of phage resistance. Proceedings of the National Academy of Sciences. 2021 Jun 8;118(23):e2104592118. 10.1073/pnas.2104592118

35. Haines ME, Hodges FE, Nale JY, Mahony J, Van Sinderen D, Kaczorowska J et al. Analysis of selection methods to develop novel phage therapy cocktails against antimicrobial resistant clinical isolates of bacteria. Frontiers in microbiology. 2021 Mar 29;12:613529. 10.3389/fmicb.2021.613529

36. Merabishvili M, Pirnay JP, De Vos D. Guidelines to compose an ideal bacteriophage cocktail. Bacteriophage Therapy: From Lab to Clinical Practice. 2018:99–110. 10.1007/978-1-4939-7395-8_9

37. Molina F, Simancas A, Ramírez M, Tabla R, Roa I, Rebollo JE. A new pipeline for designing phage cocktails based on phage-bacteria infection networks. Frontiers in microbiology. 2021 Feb 16;12:564532. 10.3389/fmicb.2021.564532

38. Naknaen A, Samernate T, Wannasrichan W, Surachat K, Nonejuie P, Chaikeeratisak V. Combination of genetically diverse Pseudomonas phages enhances the cocktail efficiency against bacteria. Scientific Reports. 2023 Jun 1;13(1):8921. 10.1038/s41598-023-36034-2

39. Forti F, Roach DR, Cafora M, Pasini ME, Horner DS, Fiscarelli EV et al. Design of a broad-range bacteriophage cocktail that reduces Pseudomonas aeruginosa biofilms and treats acute infections in two animal models. Antimicrobial agents and chemotherapy. 2018 Jun;62(6):10–128. 10.1128/aac.02573-17

40. Chan BK, Abedon ST, Loc-Carrillo C. Phage cocktails and the future of phage therapy. Future microbiology. 2013 Jun;8(6):769–83. 10.2217/fmb.13.47

41. Gadagkar R, Gopinathan KP. Bacteriophage burst size during multiple infections. Journal of Biosciences. 1980 Sep;2:253–9. 10.1007/BF02703251

42. Abedon ST. Lysis from without. Bacteriophage. 2011 Oct;1(1):46–9.

43. Cairns BJ, Timms AR, Jansen VA, Connerton IF, Payne RJ. Quantitative models of in vitro bacteriophage–host dynamics and their application to phage therapy. PLoS Pathogens. 2009 Jan 2;5(1):e1000253. 10.1371/journal.ppat.1000253

44. Krysiak-Baltyn K, Martin GJ, Stickland AD, Scales PJ, Gras SL. Computational models of populations of bacteria and lytic phage. Critical reviews in microbiology. 2016 Nov 1;42(6):942–68. 10.3109/1040841X.2015.1114466

45. Chevallereau A, Pons BJ, van Houte S, Westra ER. Interactions between bacterial and phage communities in natural environments. Nature Reviews Microbiology. 2022 Jan;20(1):49–62. 10.1038/s41579-021-00602-y

46. Nabergoj D, Modic P, Podgornik A. Effect of bacterial growth rate on bacteriophage population growth rate. MicrobiologyOpen. 2018 Apr;7(2):e00558. 10.1002/mbo3.558

47. Wiebe KG, Cook BW, Lightly TJ, Court DA, Theriault SS. Investigation into scalable and efficient enterotoxigenic Escherichia coli bacteriophage production. Scientific Reports. 2024 Feb 13;14(1):3618. 10.1038/s41598-024-53276-w

48. Sarker SA, Sultana S, Reuteler G, Moine D, Descombes P, Charton Fet al . Oral phage therapy of acute bacterial diarrhea with two coliphage preparations: a randomized trial in children from Bangladesh. EBioMedicine. 2016 Feb 1;4:124–37. 10.1016/j.ebiom.2015.12.023

49. Jault P, Leclerc T, Jennes S, Pirnay JP, Que YA, Resch G et al. Efficacy and tolerability of a cocktail of bacteriophages to treat burn wounds infected by Pseudomonas aeruginosa (PhagoBurn): a randomised, controlled, double-blind phase 1/2 trial. The Lancet Infectious Diseases. 2019 Jan 1;19(1):35–45. 10.1016/S1473-3099(18)30482-1

50. McPartland J, Rothman-Denes LB. The tail sheath of bacteriophage N4 interacts with the Escherichia coli receptor. Journal of bacteriology. 2009 Jan 15;191(2):525–32. 10.1128/jb.01423-08

51. Choi KH, McPartland J, Kaganman I, Bowman VD, Rothman-Denes LB, Rossmann MG. Insight into DNA and protein transport in double-stranded DNA viruses: the structure of bacteriophage N4. Journal of molecular biology. 2008 May 2;378(3):726–36. 10.1016/j.jmb.2008.02.059

