## Supplemental data for "The life history traits of phages in a cocktail determine coinfection dynamics and efficacy"

### Supplementary File

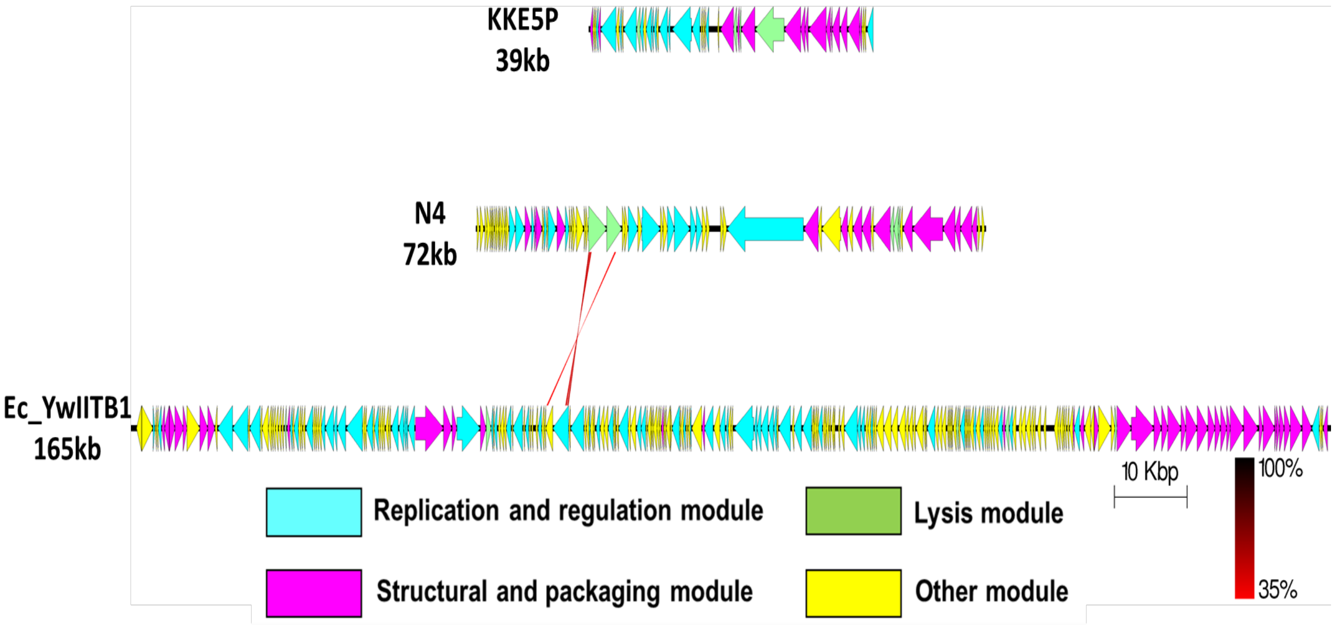


**Figure S1**: Comparison of the whole genome sequences of the phages KKE5P, N4 and Ec_YwIITB1. The colored arrows indicates CDS according to their predicted function. The homologous regions between phages are indicated by red line (Scale = base pairs).

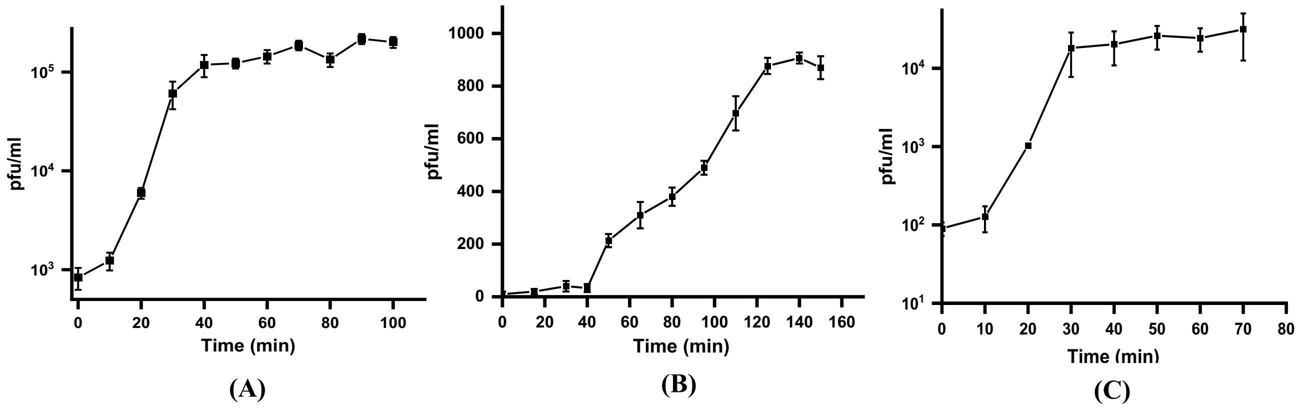


**Figure S2**. One step growth curve analysis of A) N4 Phage B) Ec_YwIITB1 Phage C) KKE5P Phage

Slope = - 0.06042 ± 0.00488

$R^2$= 0.96836

Slope = - 0.05393 ± 0.00564

$R^2$= 0.94809

Slope = - 0.08717 ± 0.01153

$R^2$= 0.93462

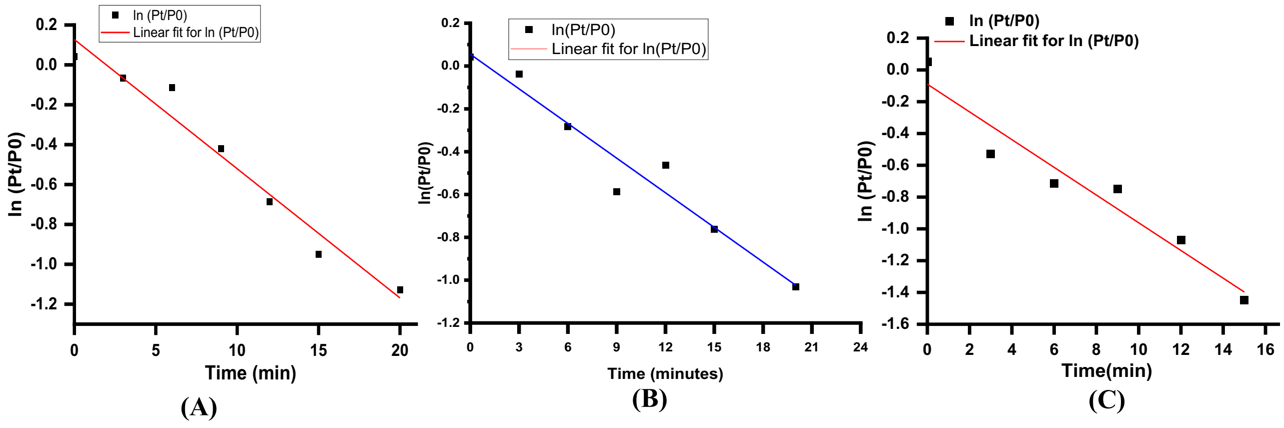


**Figure S3.** Adsorption rate analysis for A) N4 Phage B) Ec_YwIITB1 C) KKE5P Phage

Slope= - 3.28227± 0.02092

Intercept- 39.57444± 0.09204

$R^2$= 0.99976

Slope= - 3.31223± 0.03911

Intercept- 41.87416± 0.33291

$R^2$= 0.9993

Slope= -2.89875 ± 0.30779

Intercept- 40.2825 ± 2.12127

$R^2$= 0.93664

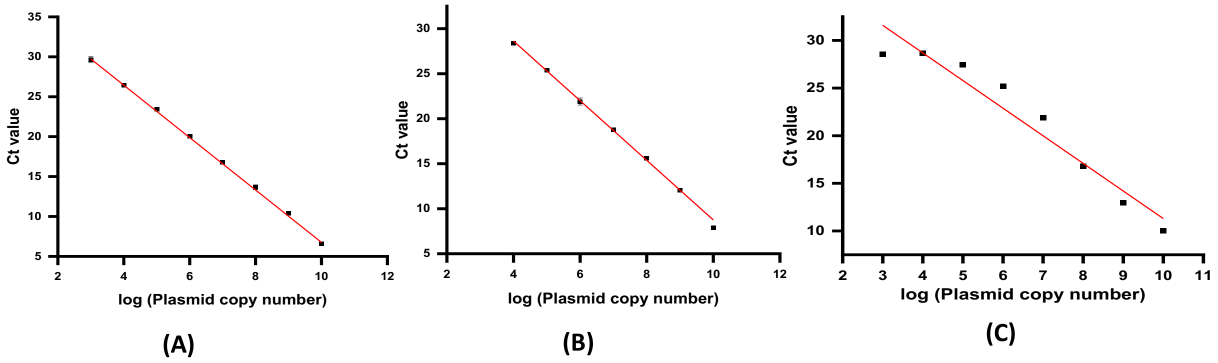


**Figure S4.** Standard curve for absolute quantification of phage gene copy number for A) N4 Phage [*g68*] B) Ec_YwIITB1 [*g22*] C) KKE5P Phage [*g10*].

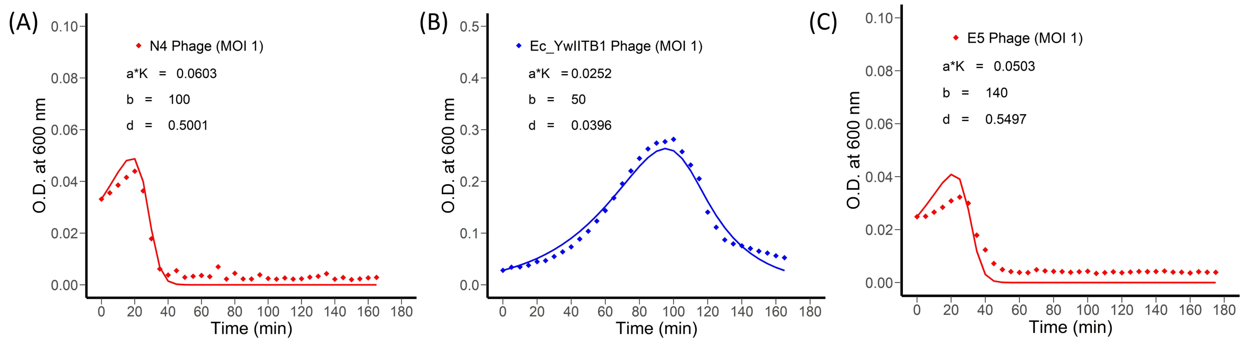


**Figure S5. Model parameter estimation after fitting with the experimental data (A) N4 phage, (B) Ec_YwIITB1 phage, and KKE5P phage.** Parameter estimation was performed using the modFit function in R.

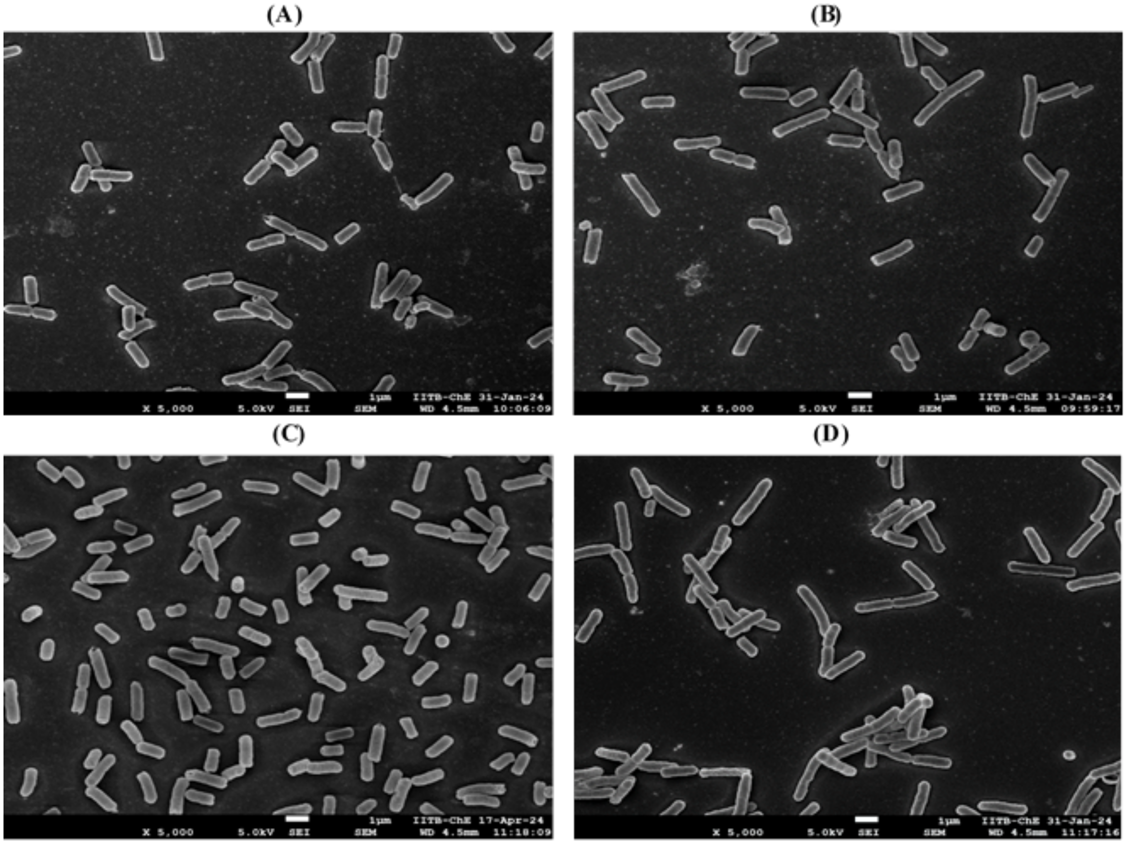


**Figure S6.** *E.coli* cell size determination by SEM analysis (A) Minimal media (B) Luria Broth (C) Terrific Broth (D) Super Broth. Scale bar- 1µm

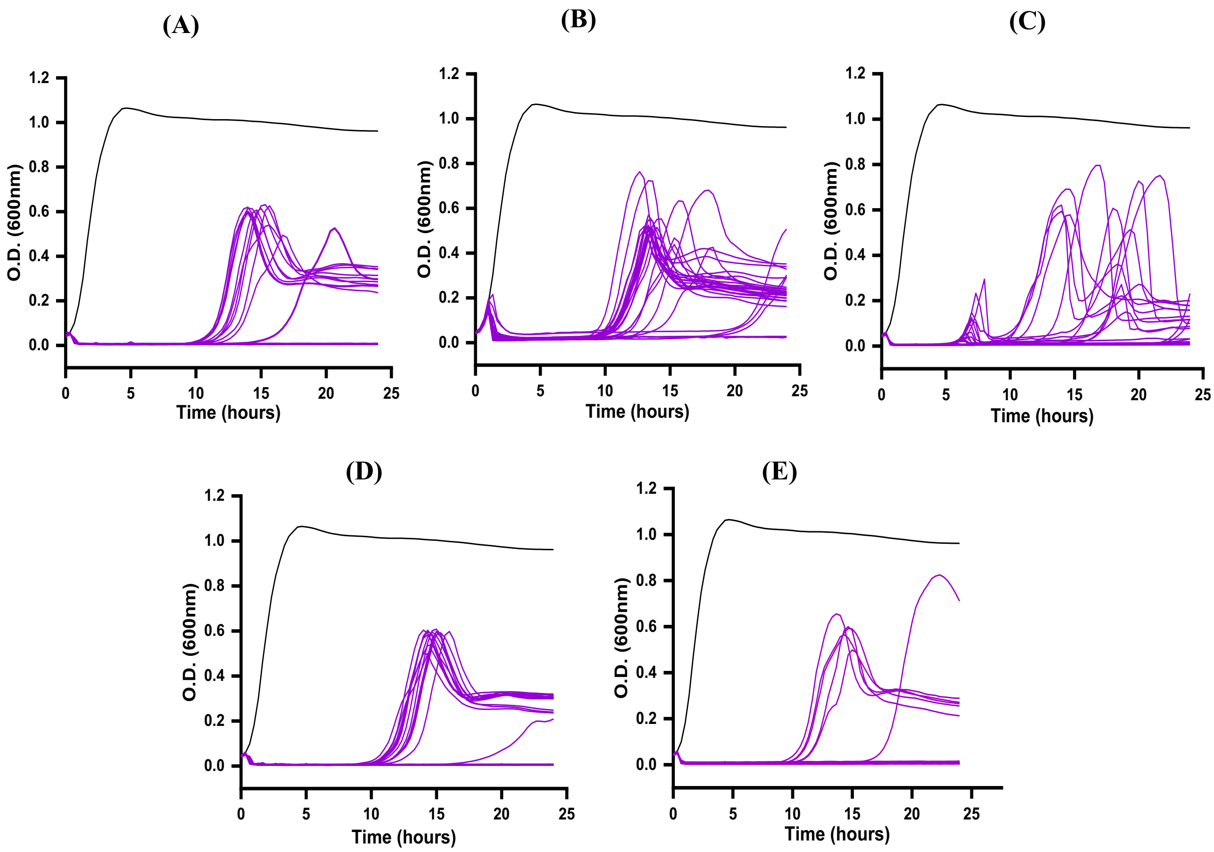


**Figure S7:** Growth trajectories of host cell infected with (A) N4 (B) Ec_YwIITB1 (C) KKE5P (D) N4+Ec_YwIITB1 (E) N4+KKE5P at MOI 1 (n=27) at 37°C. Replicates displaying regrow are termed partial resistant because of inhibited growth in comparison to uninfected while those showing no regrow are termed sensitive.

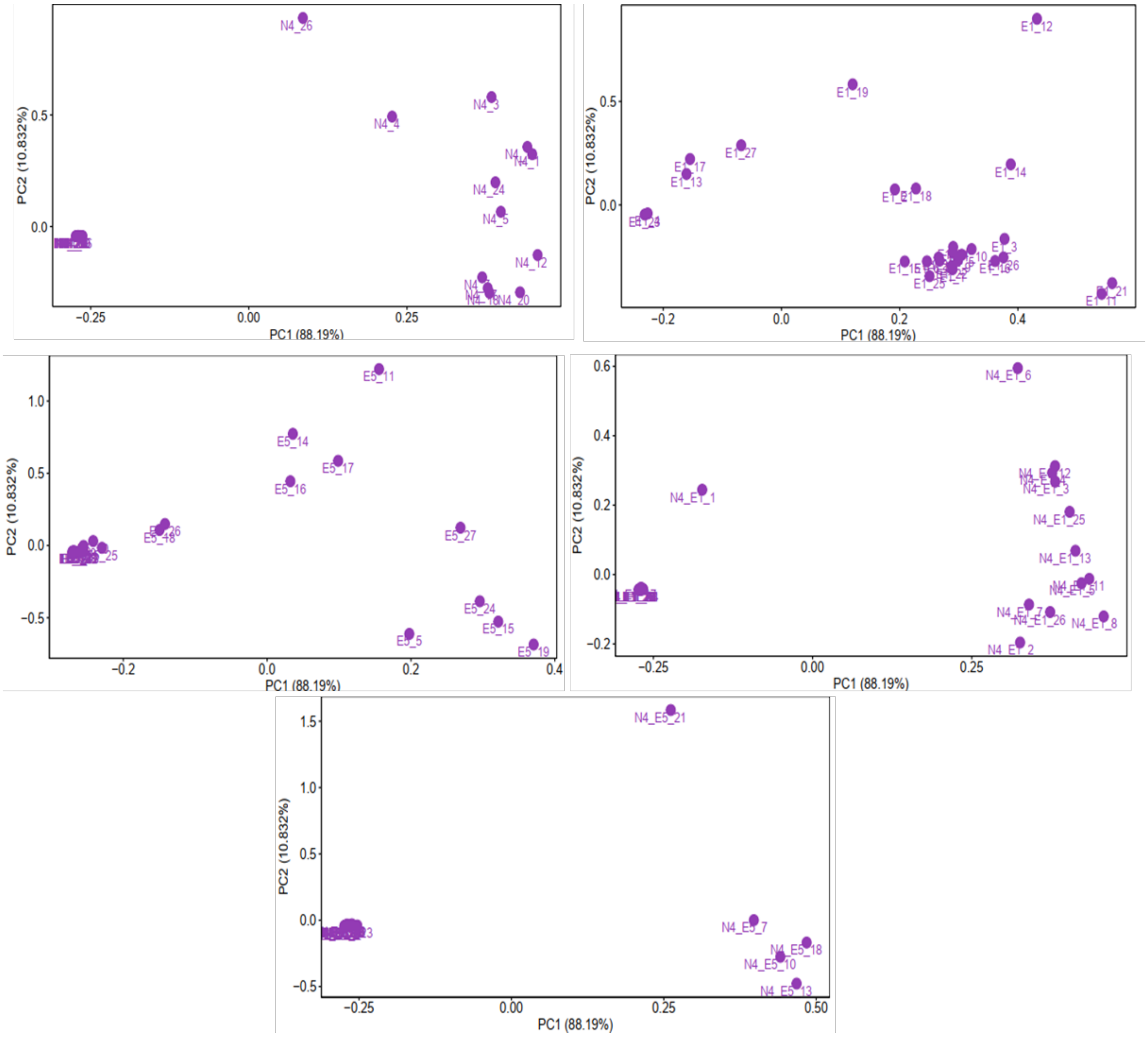


**Figure S8:** Principal Component Analysis (PCA) on the differences in OD trajectories of uninfected bacterial population and infected bacterial population in plate reader experiments of (A) N4 (B) Ec_YwIITB1 (C) KKE5P (D) N4+ Ec_YwIITB1 (E) N4+ KKE5P. PC1 vs PC2 plot represents the clustering of sensitive replicates (left side) and partial resistant replicates as per the variability of regrow among them. E1 is a short -label for phage Ec_YwIITB1.

Slope= 1.41819E9+ 2.71339E8
Intercept= 9.51032E7 ±
1.39481E8

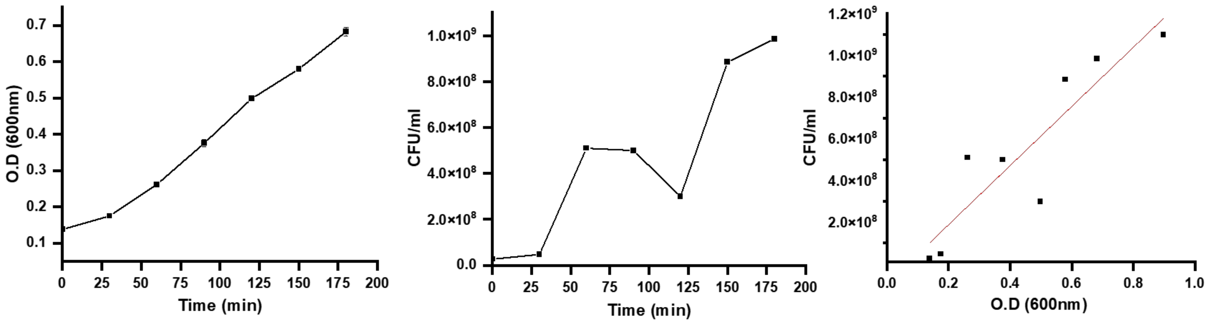


Fig S9: Standard curve for O.D. to CFU conversion

**Table S1: Primers list used for qPCR**

| Gene | Primer Sequence (5'-3') | Size (bp) |
|---|---|---|
| N4 *g68* large terminase | F: ACT GGA CAG CAA GGT GGT TT<br>R: GAG GCC AAC CGG GTA TTC AT | 105 |
| Ec_YwIITB1 *g22* | F: TCC CTT CAG AAA AAG CAG CA<br>R: TCG CCA GTT ACT TTC CAC AA | 97 |
| KKE5P *g10* | F:CCATGTAAGGACGCCAATGA<br>R:ACGTAACGAAAGAGCCGATAC | 117 |

**Table S2: Composition of various bacteriological media**

| **Media** | **Composition** |
|---|---|
| M9 Minimal Media | M9 Salt (5X)<br>$KH_2PO_4$, 15 g/L; NaCl, 5.0 g/L; $Na_2HPO_4$, 64 g/L; $NH_4Cl$, 5 g/L.<br>1X M9 Salt, 20mM glucose, 2mM magnesium sulphate<br>1mM calcium chloride |
| Luria Broth | Tryptone (casein peptone) ,10g/L; Yeast extract, 5g/L; NaCl – 5g/ L |
| Terrific Broth | Tryptone casein peptone ,12g/L; Yeast extract, 24g/L; $KH_2PO_4$ 9.4g/L; $Na_2HPO_4$ ,2.2g/L; Glycerol, 4ml/L |
| Super Broth | Tryptone (casein peptone) ,32g/L; Yeast extract, 20g/L<br>NaCl – 5g/ L |

**Table S3: Growth rate and doubling time of *E. coli* on different bacteriological media**

| Media | Growth rate | Doubling time (min) |
|---|---|---|
| M9 Minimal media | 0.00831 ± 0.0001 | 83.4 ± 0.998 |
| Luria Broth | 0.03126 ± 0.00178 | 22 ± 1.01 |
| Terrific Broth | 0.03078 ± 0.0007 | 22.5 ± 0.48 |
| Super Broth | 0.02439 ± 0.00079 | 28.41 ± 0.89 |

**Table S4. Functional and performance traits of N4, KKE5P, and Ec_YwIITB1 phages.**

| | **N4 phage** | **Ec_YwIITB1**<br><br>**(isolated and characterized in the lab)** | **KKE5P phage**<br><br>**(isolated and characterized in the lab)** |
|---|---|---|---|
| **Family** | *Podoviridae* | *Myoviridae* | *Podoviridae* |
| **Genome Size (kb)** | 72 [50] | 165 | 39 |
| **Approximate Capsid Size (diameter, nm)** | 70 | 84 | 40 |
| **Burst Size (pfu/cell)** | 135 | 35 | 197 |
| **Average Latent Period (min)** | 20 | 40 | 20 |
| **Adsorption rate (ml/min)** | $5.49 \times 10^{-11}$ | $4.9 \times 10^{-11}$ | $1.21 \times 10^{-10}$ |
| **Host range** | *E. coli* W3350 | *E. coli* MG1687<br><br>*E. coli* B40<br><br>*E. coli* W3350 | *E. coli* W3350 |
| **Recognition receptor** | Outer membrane proteins, NfrA and NfrB [51] | OmpC [predicted] | Lipopolysaccharides (LPS) [reference] |

**Table S5. Variables extracted from bacterial growth curves.**

| Variable | Description | N4 Phage | Ec_YwIITB1 | Explanation of biological correlate(s) |
|---|---|---|---|---|
| nMax | Maximum bacterial density | 0.04298 | 0.2756167 | Lysis time, burst size, attachment rate |
| tMax | Time at which bacteria reached maximum density | 20 min | 100 min | Lysis time, burst size, attachment rate |
| nExt | $OD_{600}$ reading at which bacteria went extinct | 0.0036 | 0.0755 | Relative virulence of the virus on the bacterial host |
| tExt | Time at nExt | 60 min | 140 min | Rate of viral increase |
| AUC | Area under the bacterial growth curve | 1.587 | 21.44598 | Latent period, viral fitness |

**Table S6. Estimated parameters for N4, Ec_YwIITB1, and KKE5P phage.**

| Parameter | | N4 Phage | Ec_YwIITB1 | KKE5P |
|---|---|---|---|---|
| $r$ $(min^{-1})$ (Growth rate of host bacteria) | | 0.0314 | | |
| $\alpha K$ $(min^{-1})$ (Infection rate/ adsorption rate and carrying capacity) | | 0.0603 | 0.0252 | 0.0503 |
| | Std. Error | 0.004926 | 0.005142 | 0.001577 |
| | p-value | $2.08\times10^{-13}$ | $2.83\times10^{-05}$ | <0.001 |
| $d$ $(min^{-1})$ (Death rate of host cells due to cell lysis) | | 0.5001 | 0.0396 | 0.5497 |
| | Std. Error | 0.06581 | 0.002788 | 0.01497 |
| | p-value | $1.44\times10^{-8}$ | $4.02\times10^{-15}$ | <0.001 |
| $\beta$ $(dimensionless)$ (Burst size) | | 100 | 50 | 140 |
| | Std. Error | 9.754 | 15.676520 | 1.328 |
| | p-value | $1.76\times10^{-11}$ | 0.00325 | <0.001 |

51. McPartland J, Rothman-Denes LB. The tail sheath of bacteriophage N4 interacts with the Escherichia coli receptor. Journal of bacteriology. 2009 Jan 15;191(2):525-32.

52. Choi KH, McPartland J, Kaganman I, Bowman VD, Rothman-Denes LB, Rossmann MG. Insight into DNA and protein transport in double-stranded DNA viruses: the structure of bacteriophage N4. Journal of molecular biology. 2008 May 2;378(3):726-36.
